# Rational probe design for efficient rRNA depletion and improved metatranscriptomic analysis of human microbiomes

**DOI:** 10.1101/2022.08.08.502835

**Authors:** Asako Tan, Senthil Murugapiran, Alaya Mikalauskas, Jeff Koble, Drew Kennedy, Fred Hyde, Victor Ruotti, Emily Law, Jordan Jensen, Gary P. Schroth, Jean M. Macklaim, Scott Kuersten, Brice Le François, Daryl M. Gohl

## Abstract

**Background:** The microbiota that colonize the human gut and other tissues are dynamic, varying both in composition and functional state between individuals and over time. In studying the function of the human microbiome and the mechanisms of microbiota-mediated phenotypes, gene expression measurements provide additional insights to DNA-based measurements of microbiome composition. However, efficient and unbiased removal of microbial ribosomal RNA (rRNA) presents a barrier to acquiring metatranscriptomic data, as rRNA typically accounts >90% of total RNA in microbial cells.

**Results:** Here we describe a probe set that achieves efficient enzymatic rRNA removal of complex human-associated microbial communities. We demonstrate that the custom probe set can be further refined through an iterative design process to efficiently deplete rRNA from a range of human microbiome samples, including adult and infant gut, as well as oral and vaginal communities. Using synthetic nucleic acid spike-ins, we show that the rRNA depletion process does not introduce substantial quantitative error in the resulting gene expression profiles. Successful rRNA depletion allows for efficient characterization of taxonomic and functional profiles, including during the development of the human gut microbiome.

**Conclusions:** The pan-human microbiome enzymatic rRNA depletion probes described here provide a powerful tool for studying the transcriptional dynamics and function of the human microbiome.

## Background

The microbiome plays a critical role in human health and disease [1]. Over the past decade, next-generation sequencing-based analyses have provided insights into the composition of the microbiome across body sites and life stages and have begun to uncover correlations between microbial taxa or microbial functions and disease states [2–4]. Beyond genomic analysis of microbiome composition, multi-omic data incorporate measurements of the microbiota-associated transcriptome, proteome, or metabolome to provide further insights into microbiome activity and function. Although metagenomic and metatranscriptomic profiles tend to be generally consistent, microbial functional profiles derived from DNA sequencing are more conserved across donors than transcriptional profiles, which are highly donor specific [5]. Importantly, many broadly encoded metagenomic pathways are expressed by a small number of organisms, highlighting the utility of metatranscriptomics to identify functional activities [6]. In particular, transcriptomic measurements of the human gut associated microbiome have been used to study microbial carbohydrate metabolism [7] and to provide functional information about intestinal diseases such as IBD [8] as well as mechanisms of drug metabolism [9].

Acquiring metatranscriptomic data is hindered by the fact that the vast majority of microbial-derived RNA molecules correspond to ribosomal RNA (rRNA) [10]. In eukaryotes, non-ribosomal RNA can be easily and efficiently enriched through selective reverse transcription or pull-down approaches that target the poly-A tail or using probes to specifically bind rRNA molecules prior to removal by capture or enzymatic digestion [11, 12]. Although Poly-A polymerase was first isolated from *Escherichia coli* [13, 14], bacterial mRNA transcripts are not, as a rule, poly-adenylated, and when poly-adenylation does occur it is associated with RNA degradation [15, 16]. Thus, for bacterial samples, selective enrichment of mRNA is not easily achievable and the removal of rRNA must be accomplished by other means.

While a large number of studies have developed efficient methods to deplete rRNA in individual bacterial species using probe-based capture [17], enzymatic depletion [18], or CRISPR-based methods [19, 20], depleting the diverse rRNA sequences in complex human microbiome samples that can contain hundreds of species presents a significant technical challenge. In addition, the composition of the microbiota varies substantially across body sites and throughout different life stages, further expanding the taxonomic coverage required for robust depletion of rRNA across human microbiome samples. Probe-based sequence capture methods, such as were employed with Illumina’s Ribo-Zero Gold kit can provide strong rRNA depletion across a variety of sample types, including human gut microbiome samples [21]. However, such probes are costly, difficult to manufacture, and tend to perform best with high quality RNA samples. Moreover, capture-based rRNA depletion methods can lead to inconsistent results based on operator skill. These factors led to the discontinuation of Illumina’s capture-based bacterial Ribo-Zero Gold (Epidemiology) depletion kit.

Here we describe the development of a pan-human microbiome probe set for efficient and consistent enzymatic (RNAse H) microbial rRNA depletion. Through an iterative design process, we developed probes that effectively deplete rRNA found in human oral, vaginal and adult and infant gut microbiome samples, substantially improving mapping rates to coding microbial gene databases. Using defined spike-ins, we demonstrate that the rRNA depletion process does not introduce substantial bias in the gene expression profiles. In addition, we use the resulting metatranscriptomics data to refine informatic pipelines for rRNA and host mapping and to examine gene expression and functional activity across human microbiome sites. The method described here circumvents the limitations of sequence capture methods and represents a highly effective rRNA depletion option for metatranscriptomics studies of human-associated microbial communities.

## Methods

### Samples

All samples were collected under DNA Genotek’s IRB protocol (RD-PR-00087). Human stool samples were either collected in specimen cups or in prototypes of DNA Genotek’s OMNIgene●GUT DNA & RNA (OMR-205). Unstabilized samples were returned to the lab on ice packs and either extracted within 2-3 hours or stored at −80°C until extraction. OMR-205 stabilized samples were stored at room temperature and RNA was extracted within 10-14 days of sample collection. Vaginal and oral microbiome samples were collected in DNA Genotek’s OMNIgene^®^●VAGINAL (OMR-130, vaginal microbiome sampling kit) and OMNIgene^®^●ORAL (OMR-120, tongue microbiome sampling kit) following the device IFU. Samples were stored at room temperature until further processing. Prior to extraction, collected samples were treated with Proteinase K for 1 hour at 50°C as per DNA Genotek’s instructions and concentrated down to 200-250 μl using Eppendorf’s Vacufuge Plus (centrifugation was performed at 30°C for 45-60 minutes) for optimal extraction yields.

The pooled RNA samples were generated by mixing total RNA extracted from human cells and bacterial cultures. *Francisella philomiragia* (ATCC 25017), *Escherichia coli* (ATCC 13706), *Pseudomonas aeruginosa* (ATCC 10145), *Staphylococcus aureus* (ATCC 25923), *Moraxella catarrhalis* (ATCC 25238), *Klebsiella pneumonia* (ATCC 13883), *Micrococcus luteus* (ATCC 4698), *Yersinia enterocolitica* (ATCC 9610), *Bacillus subtilis* and *Lactobacillus crispatus* (ATCC 33820) were grown overnight in their recommended culture media. Cells were pelleted, washed once with ddH2O and frozen until extraction. RNA pool 1 consists of 30% human THP-1 RNA (ATCC^®^ TIB-202™) and 10% total RNA from each the following bacterial species: *B. subtilis, E. coli, F philomiragia, L. crispatus, P. aeruginosa, S. aureus and Y. enterocolitica*. RNA pool 2 consists of equal amounts (9.09%, by ng of RNA) of total THP-1 human RNA, *F. philomiragia, E. coli, P. aeruginosa, S. aureus, M. catarrhalis, K. pneumonia, M. luteus, Y. enterocolitica, B. subtilis and L. crispatus* total RNA. Gut and skin intact cell mock communities were purchased from ATCC (cat# MSA-2006, and MSA-2005)

### RNA extractions, RNA QC and RT-qPCR

Human total RNA was extracted from THP-1 human cells using TRIzol (ThermoFisher, cat# 15596026) as per manufacturer instructions. Bacterial RNA was extracted from intact cell mock communities (ATCC, MSA-2006, and MSA-2005), bacterial cultures as well as stool, vaginal and oral microbiome samples using Qiagen’s RNeasy^®^ PowerMicrobiome^®^ Kit (cat# 26000-50), according to the manufacturer’s instructions. For raw stool samples, 50-100 mg was used as input, while for oral and vaginal samples 200-250 μl of pre-concentrated OMNIgene^®^ sample was used as input. Bead beating was performed in the presence of phenol-chloroform-isoamyl alcohol (Sigma-Aldrich, cat# 77617) and 2-mercaptoethanol (Sigma-Aldrich, cat# M6250). DNase treatment was performed on-column and total RNA samples were eluted in 100 μl nuclease-free water, then quantified using the Quant-iT™ RiboGreen™ RNA Assay Kit (ThermoFisher,cat# R11490). RNA quality and integrity of samples was checked on the Agilent 4200 TapeStation System using RNA ScreenTape or High Sensitivity RNA ScreenTape (Agilent, cat# 5067-5576 and 5067-5579). RNA integrity numbers (RINs) were highly variable across body sites/sample types and ranged from 2.0 to 9.0. Representative traces for each sample type are shown in Figure S1.

An RT-qPCR approach was used to screen the relative abundance of *Lactobacillus* in vaginal samples and control samples (pure *Lactobacillus crispatus* RNA and RNA pool 2). 50-80 ng of control or RNA extracted from OMNIgene●VAGINAL collected samples was reverse transcribed using Superscript II (ThermoFisher, cat# 18064022) or Superscript III (ThermoFisher, cat# 28025013) and random hexamers (ThermoFisher, cat# N8080127). Total and *Lactobacillus* 16S rRNA levels were quantified by qPCR using universal bacterial 16S primers (Fwd 5’-ATTACCGCGGCTGCTGG-3’; Rev 5’-CCTACGGGAGGCAGCAG-3’) and *Lactobacillus-genus* primers (Fwd 5’-ATGGAAGAACACCAGTGGCG-3’; Rev 5’-CAGCACTGAGAGGCGGAAAC-3’). 5 μl of diluted cDNA was then used as a template in qPCR reactions containing 1 μM Syto 9, 1.5 mM MgCl_2_ and 0.1 μg/μl BSA. Real-time PCR amplification was performed on the Corbett Rotor-Gene 6000 (discontinued) using the following conditions: 95°C for 2 minutes followed by 35 cycles consisting of 95°C for 30 seconds, 50°C or 55°C for 20 seconds and 72°C for 20 seconds for the *Lactobacillus-specific* primers and universal 16S primers respectively. PCR amplification was followed by an incubation at 72°C for 1 minute 30 seconds and melting (72°C to 95°C in 1°C increments). Relative abundance of *Lactobacillus* was determined by calculating ΔCt between Lactobacillus 16S and total 16S.

### Human microbiome probe pool design

Total RNA from gut microbiome samples of 9 donors [22] was processed in triplicate with the Ribo-Zero Plus rRNA Depletion Kit (using DP1 probes), converted into RNAseq libraries using the TruSeq Stranded Total RNAseq kit and sequenced on a NextSeq (PE 76), producing between 11 to 36 million reads per sample. The FASTQ [23] files from each donor were then aligned to the SILVA (v119) [24] database using SortMeRNA [25] to identify the sequences of rRNA to target for depletion. Any sequence regions that align in close proximity (1-3 nt) were merged and sorted by coverage depth and then filtered to remove any with less than 500x coverage. These regions were typically less than 200 nt in length representing rRNA segments that are not depleted by DP1 probes (Figure S3B). The top 50 most abundant regions were collected from each sample (donor) and combined to create a list of abundant regions. Any regions that overlapped were then merged and the list converted into a FASTA file. To identify and remove redundancies, a pairwise alignment of each region was performed and any regions that demonstrate equal to or greater than 80% identity were flagged and only one region was chosen for probe design (Figure S3C). The Ribo-Zero Plus probes (DP1) were then aligned to the selected, non-redundant regions and any regions where the probes were aligned at equal to or greater than 80% identity were eliminated. The remaining regions were collected, probe locations were established and antisense probe sequences were created. In addition, both HMv1 and HmV2 also include probes that were designed directly against the rRNA sequences from all 38 species present in the ATCC mock community samples (MSA-2002, MSA-2005 & MSA-2006) as well as *E. coli* and *B. subtilis*. The commercial product is called the Illumina^®^ Stranded Total RNA Prep with Ligation, Ribo-Zero Plus Microbiome and the probe pool equivalent to HMv2 is now referred to as DPM.

### RNAseq library preparation and sequencing

80 to 500 ng total RNA was used as input for rRNA depletion using either Illumina’s Ribo-Zero Gold rRNA Removal Epidemiology kit (discontinued) or Illumina’s Ribo-Zero Plus rRNA depletion kit (Illumina, cat# 20037135). For Ribo-Zero Plus depletion reactions, total RNA was mixed with 1 μl of DP1 (standard probe set) in the presence or absence of 1 to 1.5 μl of human microbiome probe pool (HMv1 or HMv2). Probes were hybridized and rRNA depleted as per manufacturer’s instructions. For undepleted samples, 10-20 ng total RNA was used as template for library preparation. RNAseq libraries were prepared using Illumina’s TruSeq Stranded mRNA Library Prep Kit (cat# 20020595) or Illumina’s Stranded Total RNA Prep Ligation Kit (cat# 20040529). Final libraries were quantified with the Quant-iT™ PicoGreen™ dsDNA Assay (ThermoFisher, cat# P7589) or the Qubit dsDNA HS Assay (ThermoFisher, cat# Q32851). Final library size was assessed by running libraries on the Agilent 4200 TapeStation System using D1000 ScreenTapes (Agilent, cat# 5067-5582) or alternatively, on Bioanalyzer High Sensitivity Chip (cat# 5067-4626). Individual libraries were then normalized, pooled, and sequenced on Illumina platforms (MiSeq, NextSeq or NovaSeq, see Supplemental Data File 1).

### ERCC spike-in experiment

To assess the performance and consistency of rRNA depletion/library preparation in complex microbiome samples, we designed an experiment where the ERCC RNA Spike-In Mix (ThermoFisher, cat# 4456740) was added to RNA samples (total stool RNA from 3 donors and RNA pool 2) either before or after rRNA depletion with Ribo-Zero Plus. The ERCC Mix contains 92 artificial transcripts of varying sizes that can be used to assess sensitivity and potential bias introduced in NGS workflows. Briefly, 1 μl of a 1/200 dilution of the ERCC RNA Spike-In Mix was mixed with 250 ng of total RNA and used as input for Ribo-Zero Plus depletion (containing the human microbiome probe pool HMv2) or added to matching rRNA-depleted samples prior to the fragmentation step of library preparation. In order to assess reproducibility, triplicates were processed for each sample and condition using Illumina’s Stranded Total RNA Prep Ligation Kit (cat# 20040529).

### Data Analysis

All statistical analyses and plotting were carried out using R [26] (version 4.0.5) and ggplot2 [27].

### Testing methods for filtering rRNA content

Validation of rRNA removal methods made use of a data set from BioProject PRJNA295252 [28].

Reads matching human genome were removed from the FASTQ files related to BioProject PRJNA295252 using bowtie2 [29] and hg37 [30]. To test the efficacy of various rRNA removal tools and databases, we tested bowtie2 [29] (version 2.4.2), SortMeRNA [25] (version 4.2.0) and three tools from the BBTools [31] (version 38.90) package (bbduk, bbsplit and seal), combined with four databases: (ar: SILVA SSU/LSU NR99[24]; art: ar + tRNA RFAM clans; ds: SortMeRNA[25] (v4.3) default database; ss: SortMeRNA[25] (v4.3) sensitive database). In order to conserve time and memory requirements, the data were subsampled for bowtie2 (2.5% or 25%), bbsplit, SortMeRNA [25] (2.5%), seal (25%) and bbduk (25% or 100%). For cmscan [32] (Infernal v1.1.4) and UProC [33] (v1.2.0), reads were merged using bbmerge-auto.sh from BBTools [31] (version 38.90), converted to FASTA using Seqtk (https://github.com/lh3/seqtk) and dereplicated using vsearch, then 1000 sequences were randomly subsampled using Seqtk and analyzed using UProC [33] by searching against the KEGG Orthologs database [34] (kegg_20140317.uprocdb.gz) and cmscan [32] by searching against the RFAM14.4 database [35].

### Host and rRNA removal

Adapter sequences were removed from the paired-end reads using Trimmomatic-0.38 [36] with a sliding window of ‘4:20’ and minimum length cut-off of 50 bases. Host and rRNA sequences were filtered using BBDuk [31] (BBTools version 38.90) using GRCh38 UCSC human genome and 5S Ribosomal RNA reference and the following categories subset from the art database: BacSSU, BacLSU, ArcSSU, ArcLSU, EukSSU, EukLSU and RFAM, for easier identification of various non-coding RNA targets. Reads that did not map to any of these databases were then processed for downstream analyses.

### Annotation of RNASeq reads

The filtered (non-rRNA and non-host) reads were annotated for taxonomy either using Kaiju [37] (version 1.7.0) by searching against the included proGenomes database or using SAMSA2 [38] (version 2) diamond search against the included RefSeq database. Functional annotation was carried out using SAMSA2 [37] (version 2) diamond search against the RefSeq functional genes and the SEED (2017) diamond database.

## Results

### Assessment of informatics tools for robust rRNA alignment and filtering

Determining the fraction of reads in a sample mapping to rRNA is a prerequisite to being able to accurately assess rRNA depletion efficacy. Thus, we first assessed the ability of five different mapping tools (bowtie2, bbduk, seal and bbsplit) [29, 31] together with four rRNA databases (SortMeRNA 4.3 (smr), bowtie2 (global and local modes, bow and bowl respectively), bbduk (duk), seal (sea) and bbsplit) [29, 31] together with four rRNA databases (SILVA[24] SSU/LSU NR99 (ar), SILVA SSU/LSU NR99 + tRNA RFAM clans (art), SortMeRNA [25] (4.3) default database. (ds), and SortMeRNA [25] (4.3) sensitive database (ss)) to reliably classify rRNA reads in depleted and undepleted samples. We examined data from a set of ten published samples [28], seven of which had high levels of rRNA and three of which had low levels of rRNA (Figure 1, Figure S2). bbduk, seal, and bbsplit were the fastest tools in our assessment and considerably faster than bowtie2 and sortmeRNA (data not shown). We detected substantial differences in performance across tools, with several of them (bowtie2, bowtie2 ran in local mode) showing elevated false negative rates and misclassifying rRNA as mRNA in high rRNA samples. Other tools (SortMeRNA) had elevated false positive rates, misclassifying mRNA as rRNA in low rRNA samples (Figure 1, Figure S2). In addition, bbduk had better efficiency in classifying mRNA and ncRNA correctly than seal and bbsplit based on uproc and cmscan results (Figure 1, Figure S2). Thus, we chose to use bbduk with SILVA SSU/LSU NR99 version 138.1 + tRNA RFAM clans for assessment of rRNA content in subsequent analyses.

**Figure 1.**
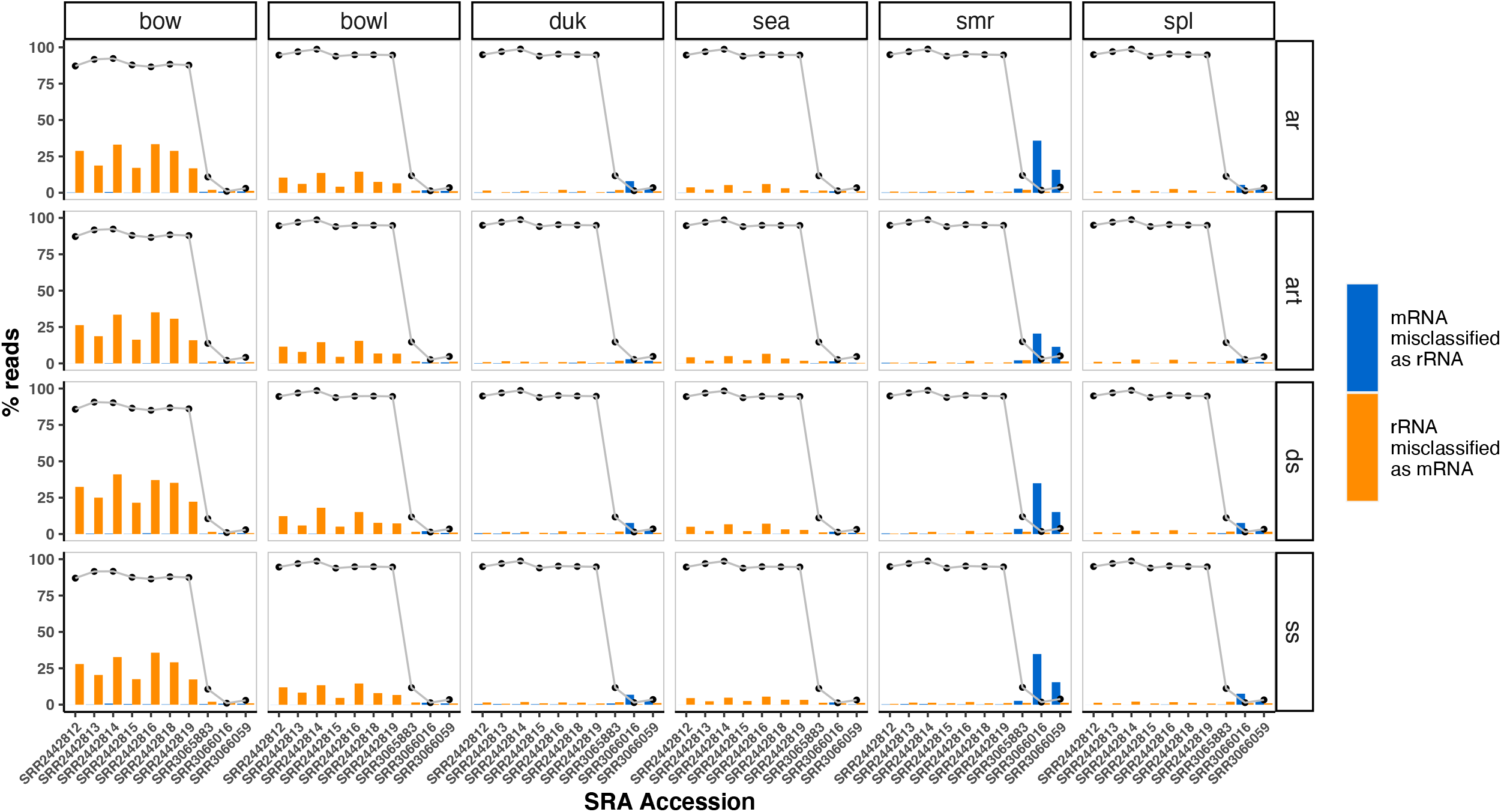
Comparison of mapping tools for classifying rRNA reads from metatranscriptomes. Bar plots showing fractions classified as either rRNA or ‘clean’ (mRNA) by the various tools (bow: Bowtie2; bowl: Bowtie2 local mode; duk: BBDuk; sea: Seal; smr: SortMeRNA: spl: bbsplit) searched against four different databases (ar: SILVA SSU/LSU NR99; art: ar + tRNA RFAM clans; ds: SortMeRNA (4.3) default database; ss: SortMeRNA (4.3) sensitive database). The line plots represent the percentage of rRNA reads in the sample identified by each method.

### Standard enzymatic Ribo-Zero Plus workflow does not efficiently deplete rRNA from complex human microbiome samples

The discontinued Illumina Ribo-Zero Gold (Epidemiology) kits used a probe-based hybridization approach to capture and deplete human and bacterial rRNAs (Figure 2A). Ribo-Zero Gold (Epidemiology) was able to substantially deplete rRNA from complex human microbiome samples such as stool, as well as from a defined pooled microbial RNA sample made up of a mix of human and bacterial total RNA (Figure 2B - see Materials and Methods for pool composition). The percentage of bacterial rRNA reads in Ribo-Zero Epidemiology samples was lower than <5% suggesting that most of the reads in these samples corresponded to bacterial and/or human mRNA. In comparison, the current Ribo-Zero Plus kit, which relies on probe hybridization followed by RNAse H enzymatic depletion (Figure 2C) performed well with the RNA pool sample but failed to substantially deplete stool samples (Figure 2D). Between 65-85% of the sequencing reads from Ribo-Zero Plus treated samples corresponded to bacterial rRNA (Figure 2D), indicating poor overall depletion efficiency for stool samples. Depletion was more successful for the less complex RNA pool sample, where only <7% of the sequencing reads mapped to the SILVA rRNA database [24]. The fact that more complex microbiome samples were less effectively depleted suggested that the standard probe content in Ribo-Zero Plus (probes targeting *E. coli* and *B. subtilis* rRNA) provides insufficient coverage to target the greater diversity of rRNA found in human gut microbes for enzymatic degradation.

**Figure 2.**
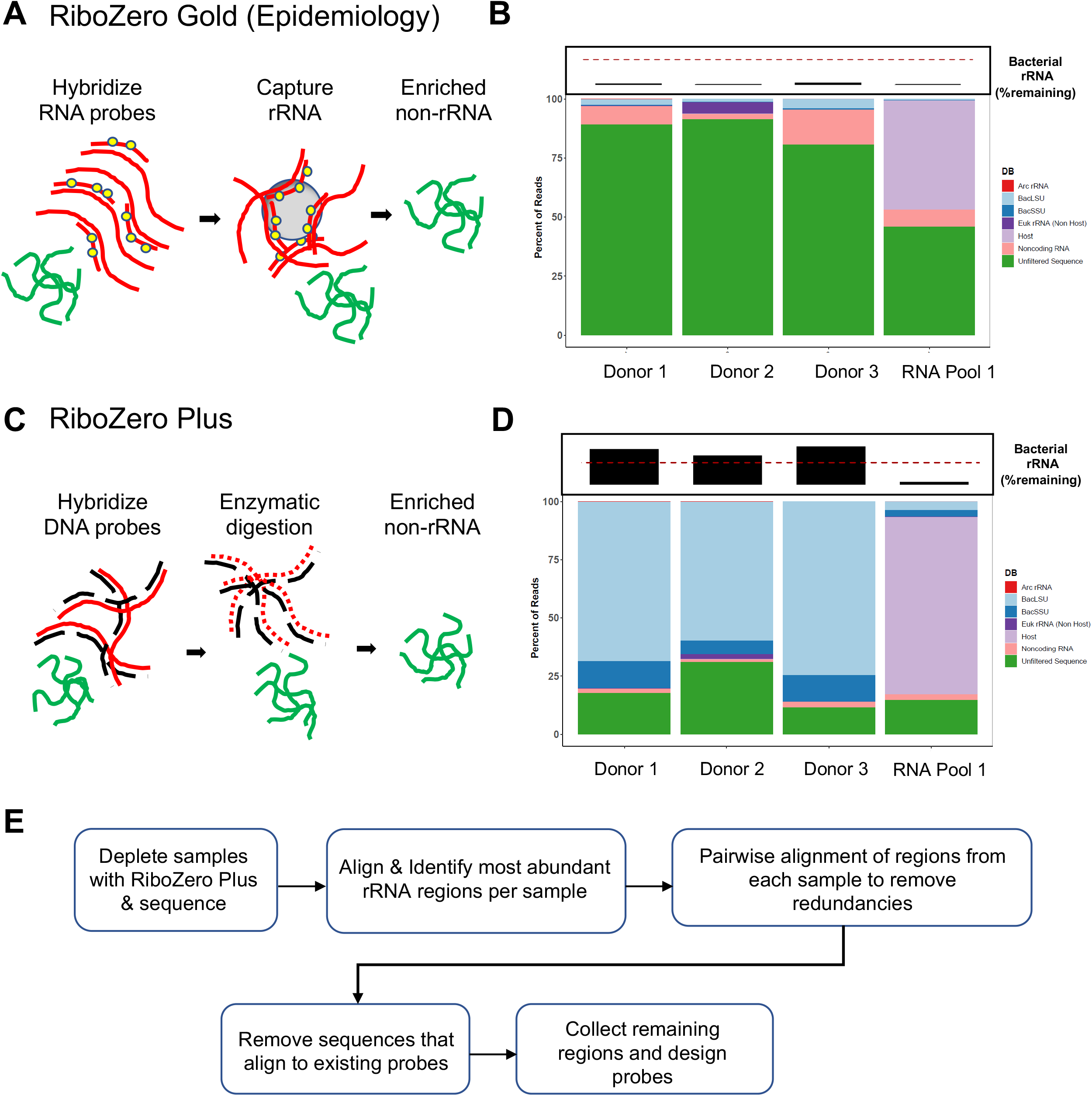
Comparison of Ribo-Zero Gold (Epidemiology) and Ribo-Zero Plus performance for rRNA removal from total RNA extracted from stool samples versus a synthetic RNA pool containing a mix of bacterial and human RNA. A) Overview of the Ribo-Zero Gold (Epidemiology) workflow that captures rRNA using biotin labeled anti-sense RNA probes captured by streptavidin magnetic beads for removal of rRNA from the sample. B) Percentage of reads mapping to eukaryotic and prokaryotic rRNAs vs coding sequences following depletion of adults stool sample and RNA standard with Ribo-Zero Gold (Epidemiology). Boxed bar plots on top represent remaining bacterial rRNA (LSU & SSU) in each sample with a dashed line at the 50% mark. C) Overview of the Ribo-Zero Plus method where anti-sense DNA oligonucleotides are hybridized to rRNAs in the sample prior to enzymatic digestion of the rRNA:DNA duplexes with RNase H. D) Percentage of reads mapping to eukaryotic and prokaryotic rRNAs vs coding sequences following depletion of adults stool sample and RNA standard with Ribo-Zero Plus. Boxed bar plots on top represent remaining bacterial rRNA (LSU & SSU) in each sample with a dashed line at the 50% mark. E) Simplified diagram of the iterative probe design process, starting from raw sequencing data of samples depleted with the standard Ribo-Zero Plus probe set (DP1). A more detailed description can be found in Supplemental Figure S3 and in Materials and Methods.

### Iterative design of a microbiome depletion oligo pool enables robust rRNA depletion from human stool samples

To improve enzymatic depletion using the Ribo-Zero Plus kit we used an iterative design process to generate additional probes specifically targeting human gut microbiome samples (Figure 2E, Figure S3). We used sequencing data from stool samples depleted with the standard Ribo-Zero Plus kit and identified the most abundant rRNA sequences that were not effectively depleted across 9 adult healthy stool RNA samples. After eliminating redundancy within these sequences and redundancy with the initial Ribo-Zero Plus probe set (DP1), we designed and synthesized a novel probe set (human microbiome pool, HMv1). Addition of HMv1 probes to Ribo-Zero Plus depletion reactions dramatically improved bacterial rRNA depletion of a group of ten test stool samples that were poorly depleted by DP1 alone (Figure 3A). The percentage of bacterial rRNA reads was >70% on average for samples depleted with DP1, while the percentage of rRNA reads was <17% on average in samples depleted with the supplemental HMv1 oligo pool (Figure 3A). In contrast, for undepleted stool samples bacterial rRNA represented over 98% of all reads. The abundant taxa (Figure 3B) and functional profiles (Figure S4) detected in the metatranscriptomic data resulting from HMv1 depletion are consistent with published human gut microbiome data, with a high abundance of taxa such as *Faecalibacterium, Lachnospiraceae, and Clostidium*. Addition of the HMv1 probes improved the number of mapped non-rRNA reads and increased the number of taxa and functional features observed (Figure 3C).

**Figure 3.**
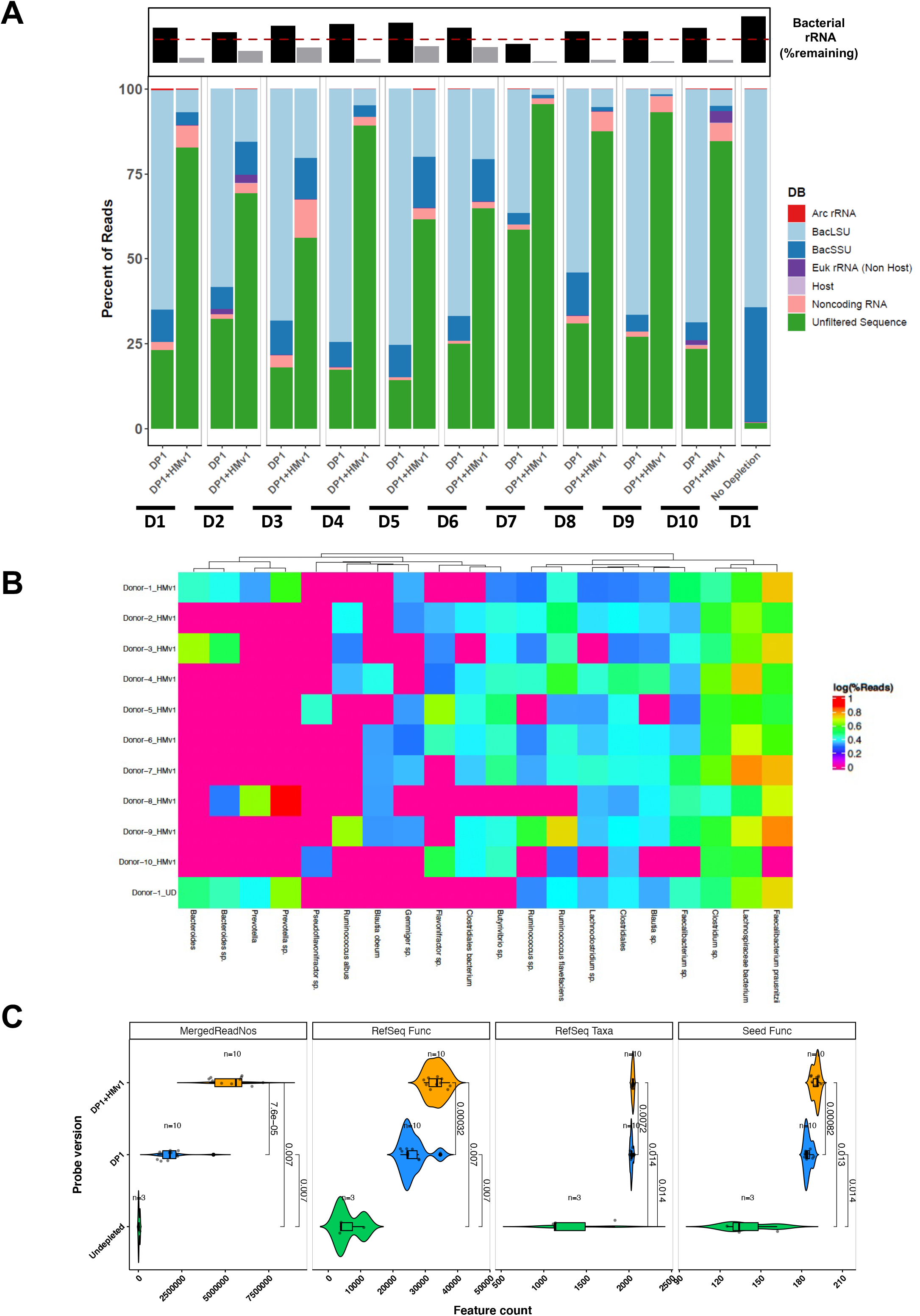
The Pan-Human Microbiome probe pool, HMv1, effectively depletes rRNA in stool samples from healthy adult donors to generate metatranscriptomic profiles. A) Percentage of reads mapping to eukaryotic and prokaryotic rRNAs vs coding sequences in stool samples depleted with Ribo-Zero Plus standard probes (DP1) vs combination of DP1 and HMv1 probes in stool RNA samples from 10 healthy adult donors. Boxed bar plot on top represents remaining bacterial rRNA (LSU & SSU) for either DP1-depleted samples (black bars) or HMv1-depleted samples (grey bars) with a dashed line at the 50% mark. B) Taxonomic heatmap showing the 20 most abundant taxa for each sample based on metatranscriptomic analysis of adult stool sample. C) Number of reads, taxonomic and functional features detected in undepleted, DP1-depleted, or HMv1-depleted stool samples.

### Pan-human microbiome oligo pool allows robust rRNA depletion of diverse human microbiome samples

Microbial community structure varies widely between donors and across body sites [39]. To assess the performance of the HMv1 probe pool across a variety of human microbiome sample types, we first depleted mock microbial communities using Ribo-Zero Plus. The Ribo-Zero Epidemiology kit was used as a control in these experiments and showed efficient depletion of rRNA in mock skin and to a lesser extent gut mock communities (Figure S5). However, variability was high between replicates especially for the gut mock. This is a well-known issue with probe capture-based workflows, where operator-dependent variables such as the amount of bead carry-over can greatly impact rRNA depletion. In contrast, Ribo-Zero Plus depletions were more reproducible (Figure S5). Interestingly, the standard Ribo-Zero Plus probe set, efficiently depleted rRNA from the gut mock community (~15% of reads mapping to rRNA) but did not perform as well with the skin mock (~50% reads mapping to rRNA). Poor performance of the Ribo-Zero Plus kit with the lower complexity skin mock community was unexpected, but further analysis revealed that it was due to inability to target *Corynebacterium striatum* and *Micrococcus striatum* rRNA sequences. Addition of the HMv1 probe set to the Ribo-Zero Plus reactions greatly improved depletion efficiency of the skin communities with <2% of reads mapping to rRNA (Fig S5). This demonstrates that the HMv1 custom probe set provides coverage against rRNA sequences from these two species (see Material and Methods).

To further test the efficacy of the HMv1 probe set on human microbiome samples, we depleted human oral (tongue) and vaginal microbiome using the Ribo-Zero plus kit. The standard Ribo-Zero Plus kit was able to efficiently deplete rRNA of half of the vaginal samples V1-V3 but performed sub-optimally for samples V4-V6 (Figure 4A). These three samples depleted significantly better following addition of HMv1. Moreover, the standard Ribo-Zero Plus kit (without supplemental probes) also performed poorly for higher complexity oral microbiome samples (Figure 4B, Figure S6). Addition of HMv1 was most striking for oral microbiome samples, where bacterial rRNA reads decreased from approximately 45% to 5% on average (Figure 4B and Figure S6). qPCR and taxonomic analysis of the vaginal metatranscriptomic profiles showed that samples efficiently depleted by the standard probe set were dominated by *Lactobacillus (V2* & V3) or *Corynebacterium* (V1), while samples benefiting from addition of HMv1 had generally a more complex community structure and higher relative abundance of *Gardnerella, Lactobacillus* and *Olsenella* (Fig. 4C and Figure S6). In comparison, metatranscriptomic profiles and diversity of tongue microbiome samples were highly consistent across donors, with *Veillonella, Rothia, Streptococcus* and *Prevotella* being the most abundant genera (Figure 5D). Taken together, our data demonstrates that the custom HMv1 probe pool improved rRNA depletion of complex human microbiome samples through increased coverage of bacterial rRNA sequences found across various sites.

**Figure 4.**
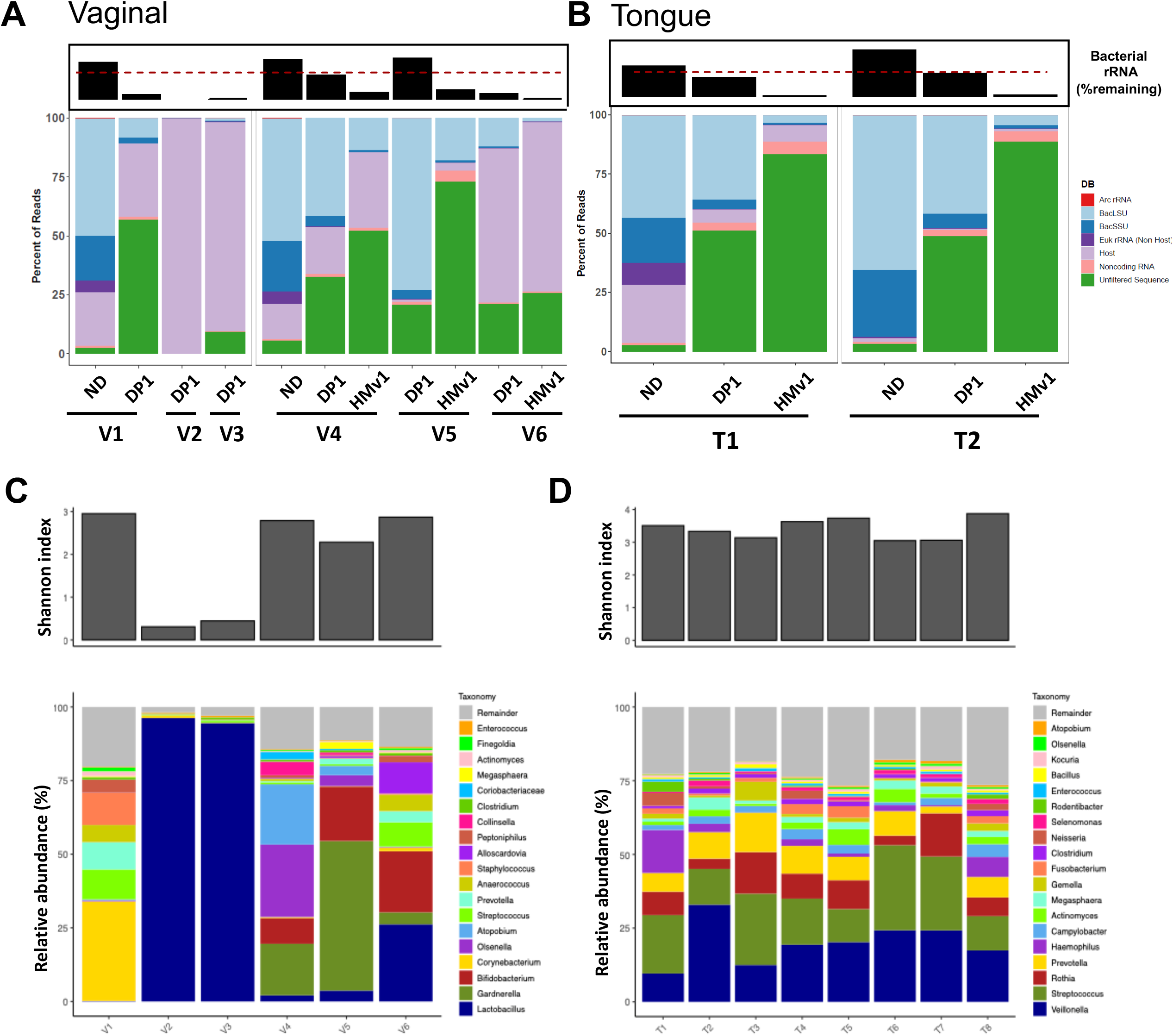
The Human microbiome pool probe set can effectively deplete rRNAs found in vaginal and oral microbiome samples to generate metatranscriptomic profiles. Percentage of reads mapping to eukaryotic and prokaryotic rRNAs vs coding sequences following depletion of vaginal and tongue samples using Ribo-Zero Plus +/− pan-microbiome HMv1 probes. A) rRNA depletion efficiency of vaginal samples using the Ribo-Zero standard probe set (DP1) +/− pan-microbial HMv1 probe set. Non-rRNA host RNA content can be prominent in vaginal samples. ND = Non depleted controls. B) rRNA depletion efficiency of representative tongue microbiome samples (T1 & T2) using the Ribo-Zero Plus standard probe set (DP1) +/− pan-microbiome HMv1 probe set. Boxed bar plots on top in panels A and B represent remaining bacterial rRNA (LSU & SSU) for indicated samples with a dashed line at the 50% mark. C) Alpha diversity (Shannon index) and metatranscriptomic taxonomic profiles for Ribo-Zero Plus (DP1 or DP1 + HMv1) depleted vaginal microbiome samples. D) Alpha diversity (Shannon index) and metatranscriptomic taxonomic profiles for Ribo-Zero Plus (DP1 + HMv1) depleted oral microbiome samples.

**Figure 5.**
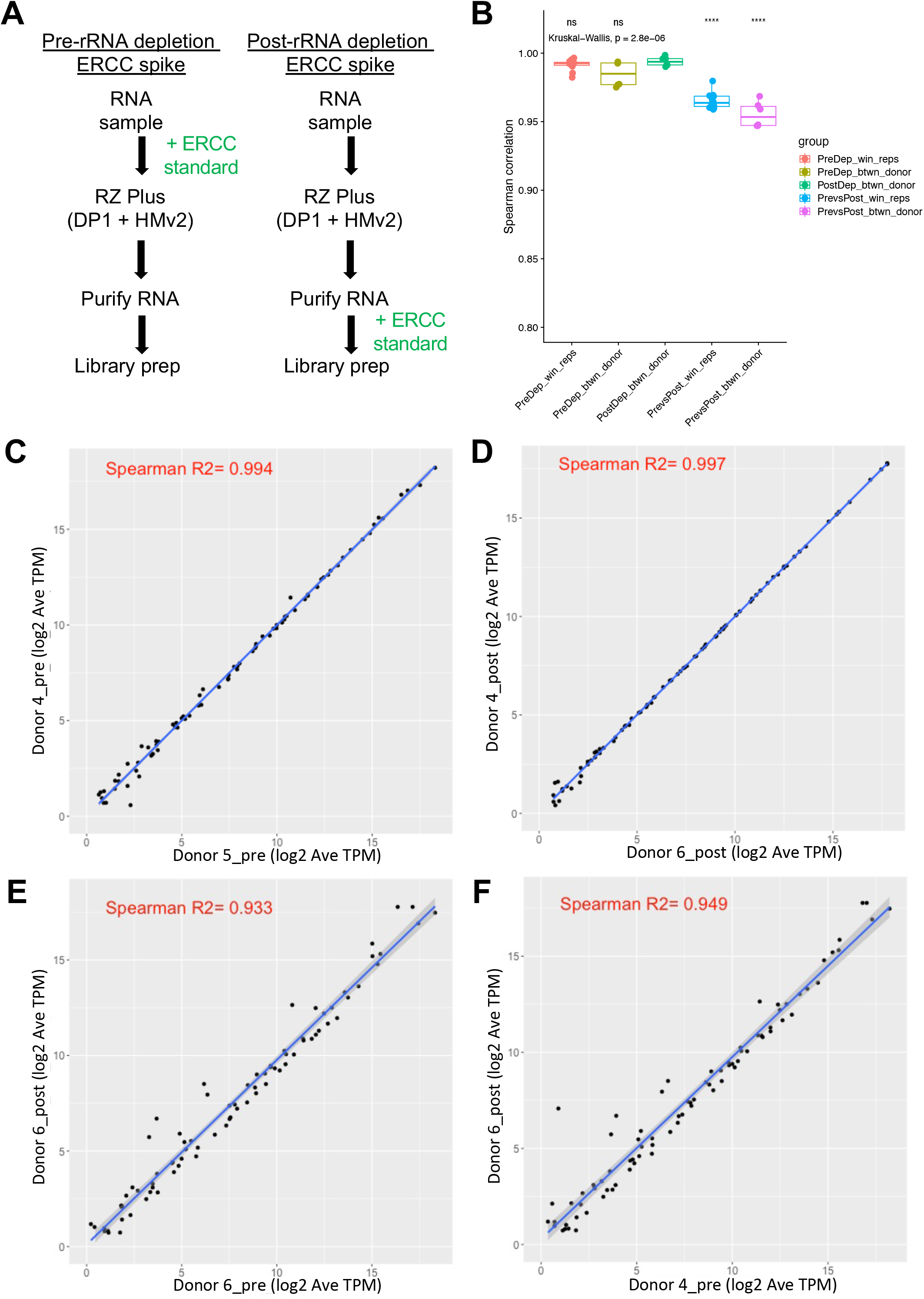
Addition of ERCC spike in controls prior to or after rRNA depletion to measure the accuracy of rRNA depletion and potential library prep bias caused by the DP1 and HMv2 probes. A) ERCC controls were spiked into the library prep either immediately before or after rRNA depletion. B) Spearman correlations between different groups tested. C-F) Representative pairwise correlations between ERCC spike-in abundance for sample pairs with the following characteristics: C) pre-depletion, different donors, D) post-depletion, different donors, E) pre- vs. post-depletion, same donor, F) pre- vs. post-depletion, different donors.

### Assessing the accuracy and reproducibility of rRNA depletion using Ribo-Zero Plus with the supplemental pan-human microbiome oligo pool

In order to assess the potential impact of the rRNA depletion process on the quantitative accuracy of the resulting metatranscriptomic measurements, we carried out an experiment where synthetic External RNA Controls Consortium (ERCC) spike-ins were added to samples pre- or post-depletion with Ribo-Zero Plus (Figure 5A). Fecal samples from 3 healthy adult donors as well as an RNA pool were tested in triplicate to assess reproducibility and accuracy of the rRNA depletion workflow. As expected efficient rRNA depletion was seen for all samples and ERCC spike-in mRNAs represented between 5-40% of total reads (Figure S7). The higher proportion of ERCC reads recovered in samples spiked post-depletion is due to underestimation of the absolute rRNA depletion that was achieved (Figure S7). Both within and between donors, the correlation between samples spiked pre-depletion and spiked post-depletion were universally high (Figure 5B). When replicates were averaged, all combinations of samples spiked pre-depletion had Spearman correlation coefficients higher than 0.95 (Figure 5B). ERCC spike-in abundances for a representative pair of pre-depletion samples are shown in Figure 5C. Likewise, samples that were spiked post-depletion also had Spearman correlation coefficients greater than 0.95 (Figure 5B, D). Comparison of pre- and post-depleted samples allow assessment of the amount of quantitative error introduced by the depletion process. Both within donor (Figure 5E) and between donor (Figure 5F) comparisons of samples spiked with ERCC spike-ins pre- and post-depletion had lower correlation coefficients than comparisons within either the pre- or post-depleted sample sets (Figure 5B). However, the correlation of ERCC spike-in abundances remained high for pre- vs. post-depletion samples. Thus, while the process of rRNA depletion introduces a detectable shift in the relative abundance of the ERCC standards, the quantitative accuracy and reproducibility of the resulting measurements remains high.

### Additional probes improve rRNA depletion of infant stool microbiome samples

Human gut microbiome profiles are known to change rapidly during the first few years of life [40]. In young infants, the gut microbiota is significantly different from adult samples and tends to be dominated by different taxa such as Bifidobacteria [41]. To determine if the Ribo-Zero Plus HMv1 probe set can efficiently remove rRNA in infant gut microbiome samples, we extracted total stool RNA collected from infants and young children aged 4 to 33-months. The HMv1 probe pool led to efficient and consistent rRNA depletion of most infant stool samples (Figure 6A and Figure S8) with <26% of reads mapping to bacterial rRNA on average. Interestingly, rRNA depletion was less efficient for a subset of donors (infants F, I, J and L) primarily in the 9 to 14-months old group. Taxonomic analysis revealed that these samples had high levels of *Bifidobacterium bifidum*. Lack of depletion suggests that the HMv1 probe set poorly targets rRNA from this species. Additional probes targeting the *Bifidobacterium bifidum* rRNA sequence were designed using our iterative process and added to HMv1 probe to create a second human microbiome pool (HMv2). To test the effectiveness of the new probe set, RNA samples from infants F, I J and L were depleted with the Ribo-Zero Plus kit using HMv1 or HMv2. The HMv2 probe set was able to improve rRNA depletion in all 4 samples compared to HMv1 (inset Figure 6A, Figure S8), with average bacterial rRNA content decreasing from 35% to 16%. The HMv2 pool was also tested on adult samples and compared to HMv1 and led to slightly better depletion efficiency in some samples (Figure S9). Unlike infant stool samples, better depletion in adult samples was not attributable to improved depletion of *Bifidobacteria*, but rather to improved rRNA coverage and depletion of unrelated taxa such as *Blautia*. This demonstrates that additional bacterial rRNA sequences can be targeted by the iterative probe design process to further improve the performance of the human microbiome probe pool. Despite marginally better rRNA depletion, HMv2 did not significantly impact the number of features detected relative to HMv1, however, the total numbers of raw reads were different in these experiments making a direct comparison difficult (Figure S10).

**Figure 6.**
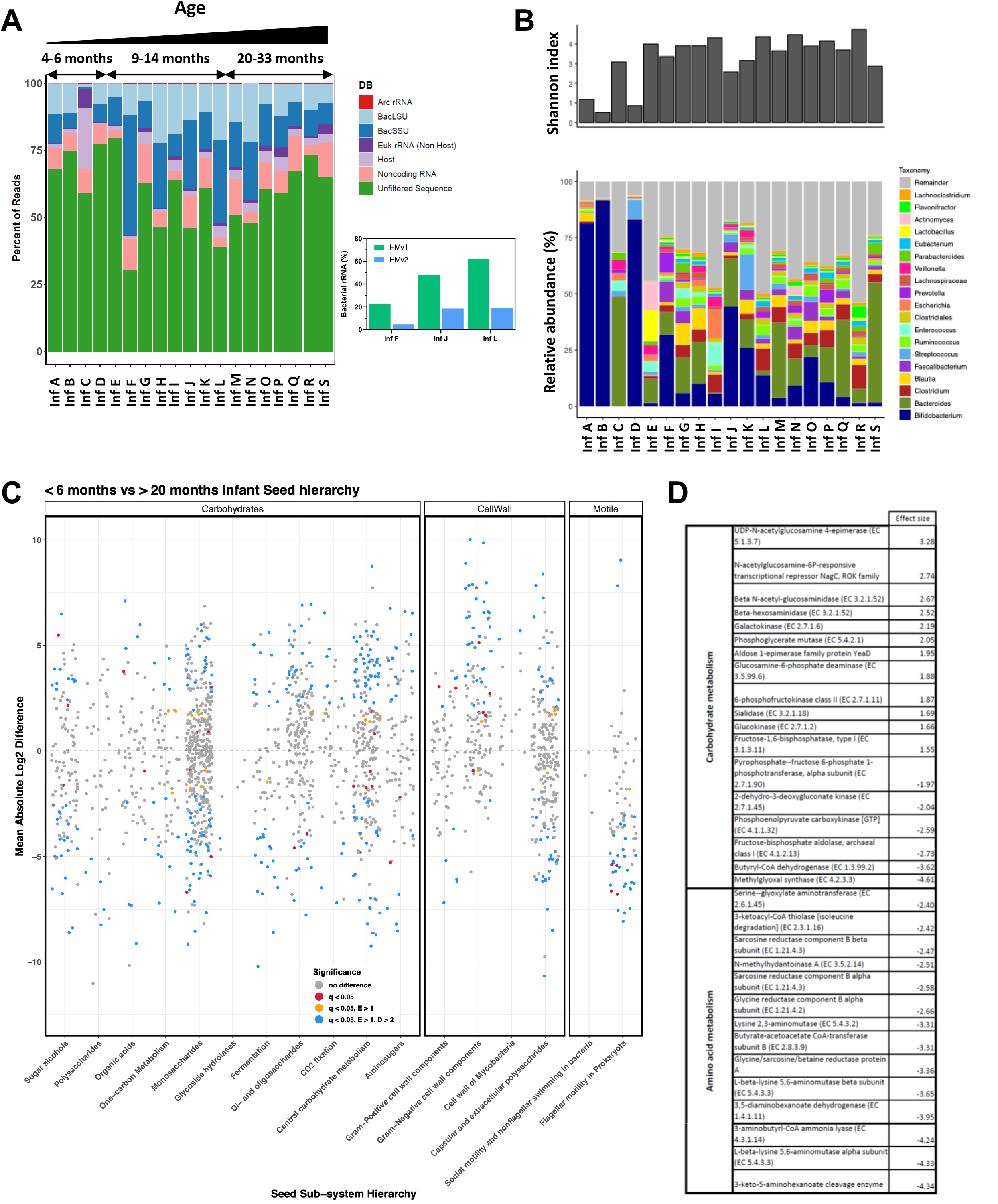
Iteration on the Human Microbiome probe pool to achieve optimal depletion of infant stool samples for metatranscriptomic analysis. A) Percentage of reads mapping to eukaryotic and prokaryotic rRNAs vs coding sequences following depletion of infant stool RNA samples 4-33 months of age using Ribo-Zero Plus and HMv1 probes (Left). Bacterial rRNA depletion (LSU & SSU) efficiency using HMv1 vs HMv2, a modified probe set supplemented with 42 additional probes targeting *Bifidobacterium bifidum* rRNA for three infant samples which had suboptimal depletion with the HMv1 probes is shown in the inset on lower right. B) Alpha diversity (Shannon index) and metatranscriptomic taxonomic profiles for HMv1 depleted stool samples from infants and children aged 4 to 33 months C) Differential gene expression in young (<6 month of age) vs older infants (>20 months) across major metabolic groups (carbohydrate metabolism, cell wall and motility). D) Table showing carbohydrate and amino acid metabolism genes with large changes in relative expression in infants (<6 months) versus young children (>20 months). Positive effect size values correlate with increased expression in infants while negative values correlate with increased expression in >20 month children.

### Stool metatranscriptomic analyses reveal significant differences in functional profiles across age groups

The effective depletion of rRNA from both infant and adult stool samples provided an opportunity to compare the taxonomic and functional profiles of these stool samples across age groups. Infants <6 months (A, B, C and D), generally had a lower Shannon index than older infants (Figure 6B – top panel) and their metatranscriptomic taxonomic profiles were largely dominated by *Bifidobacteria (*Figure 6B – bottom panel). *Bifidobacteria* relative abundance started decreasing in young children >6 months old, concomitantly with the appearance of canonical gut commensal genera such as *Faecalibacterium* and *Bacteroides* and an increase in alpha diversity (Figure 6B). Interestingly, differential abundance analysis revealed higher prevalence of *Escherichia, Veillonella, Klebsiella*, and *Shigella* species in younger infants, taxa not typically associated with healthy gut samples, as well as >20 species of the *Enterococcus* genus (Figure S11). In contrast, older infants displayed higher prevalence of several species of the *Alistipes* and *Akkermansia* genera, as well as *Mucinivorans hirudinis and Clostridium viride*. Comparison of the genes differentially expressed in infants and children of varying ages also showed significant differences, spanning many major enzymatic groups (Figure S12-13). To further understand how gut metatranscriptomic profiles evolve with age, we compared gene expression in infants (<6 months) and young children (>20 months). Infants displayed higher levels of enzymes involved in amino sugar metabolism, glycolysis, pyruvate and succinate metabolism, protein translation and cell growth (Figure 6C-D, Supplemental Data File 2). Samples from >20 month children displayed different metabolic profiles, with a large representation of genes involved in the degradation of amino acids such as glycine and lysine, butanoate fermentation (Figure 6D, Supplemental Data File 2). Genes involved in sporulation and motility were also significantly increased in >20 month old children, correlating with the appearance of spore-forming and/or motile bacteria (Figure 6B). Interestingly, both young infants and >20 month old children displayed strong upregulation of stress response pathways (cold shock proteins, chaperones, universal stress protein, two-component system), yet specific gene expression appeared to remain quite distinct in both groups (Supplemental Data File 2). We also compared gene expression in adult versus infant stool samples and observed marked differences in metabolic activity. Many enzymes of the glycolysis and the pentose phosphate pathways were increased in infants as well as downstream metabolic enzymes (such as pyruvate oxidase, pyruvate dehydrogenase, and lactate dehydrogenase) (Supplemental Data File 3). Other upregulated genes included sugar transporters (galacto-N-biose-/lacto-N-biose ABC transporter) as well as several genes associated with cell division. Stress response pathways were heavily represented in infants with >10 chaperone/cold shock proteins showing increase expression (Supplemental Data File 3). In contrast, adult samples showed little to no stress response but expressed high levels of a wide variety of sugar transporters and glycosylases (Supplemental Data File 3). Additionally, adult samples also showed high expression of genes involved in epithelium invasion/adherence, quorum sensing and biofilm formation as well as increased expression of several enzymes of the benzoate degradation and acetogenic pathways (Supplemental Data File 3).

## Discussion

Here we describe the development of probes for enzymatic rRNA depletion of human-associated microbiomes to enable metatranscriptomic analysis. First, in order to generate accuratet measures of rRNA depletion of human microbiome samples, we assessed the ability of different sequence alignment algorithms to accurately classify microbial rRNA and mRNA reads across depleted and undepleted stool samples available in public databases. Bbduk had minimal false negative and false positive mapping rates across a broad range of rRNA content and was therefore used in subsequent analyses (Figures 1, S2). Next, we used an iterative design process to design probes that effectively deplete rRNAs found in commonly studied human microbiomes (Figures 2, S3) for the Ribo-Zero Plus workflow. This enzymatic rRNA depletion approach overcomes issues associated with the cost of synthesizing sequence capture-based rRNA depletion probes and the variability and dependance of capture-based rRNA depletion on both operator skill and RNA quality.

Using the newly designed custom depletion probe set, we demonstrated robust rRNA depletion of human-associated microbiota across body sites and developmental stages, including adult and infant gut samples (Figures 3, 6, S8-S9) as well as human oral and vaginal samples (Figure 4). The human microbiome probe set was also effective at depleting rRNA from a skin mock community (Figure S5). Of all the human microbiome sites included in our study, vaginal samples and mock communities were the only ones that did not always require addition of the custom human microbiome probe set. *Lactobacillus-dominated* samples were well depleted by the standard Ribo-Zero Plus rRNA depletion probes while more complex samples with high levels of *Gardnerella, Corynebacterium*, or *Bifidobacterium* typically associated with vaginosis [42, 43] required additional probes for optimal depletion (Figure 4). This suggests that in general increased bacterial diversity correlates with increased presence of unique rRNA sequences in the sample, which in turn require additional coverage for successful depletion using RNAse H-based depletion strategies. However, there were two notable exceptions to this rule: V1, which was dominated by *Corynebacterium, Streptococcus*, and *Prevotella* depleted well with the standard probes, while the ATCC-2005 skin mock required HMv1 for optimal results. Surprisingly, both samples had high levels of *Corynebacterium*, albeit not the same species or strains. Nevertheless, we also showed that the probe set can be further refined if needed to add sequence coverage for additional targets in cases where depletion is sub-optimal. For certain infant samples, addition of probes targeting *Bifidobacteria* improved rRNA depletion (Figures 6, S8). Interestingly, the same was true for adult samples that lacked *Bifidobacteria* (Figure S6). Both HMv1 and HMv2 were designed to target abundant stool rRNA sequences, yet unexpectedly these probe sets show efficient depletion of rRNA in microbiome samples with completely different community structure (oral and vaginal). This demonstrates that pan microbiome rRNA depletion can be achieved through broad and strategic coverage of slightly divergent rRNA sequences. The ability to iterate and supplement the base rRNA depletion probe set may be important for microbiome studies especially non-western or non-industrialized populations where microbiome structure and diversity may vary significantly [44]. In addition, this iterative probe design approach provides a roadmap for the future development of depletion probes for other complex microbiome samples such as environmental samples or soil.

This enzymatic-based rRNA depletion method relies on hundreds of DNA probes targeting the diversity of rRNA sequences found in complex microbial community, an approach that has the potential to introduce bias and off-target effects. Therefore, we used synthetic spike-ins to assess the impact of the rRNA depletion process on the quantitative accuracy and reproducibility of gene expression measurements (Figure 5). All ERCC transcripts were detected and their relative abundances were highly reproducible across all of the conditions and samples tested. Comparison of samples spiked pre- and post-depletion indicated that there was a small, though detectable effect of rRNA depletion on ERCC transcript abundance. However, the Spearman correlations of ERCC transcript abundance between pre- and post-depleted samples were >0.95 among replicates from the same sample as well as across samples, thus the quantitative bias introduced by the rRNA depletion process is minimal and depletion is not expected to introduce significant error in the resulting microbial gene expression measurements.

To date, the microbiome field has only limited insights on gene expression patterns in the gut microbiome [6, 45], and only a few comparative metatranscriptomic analyses have been performed. Abu-Ali *et al*. looked at the transcriptomics profiles of 372 healthy adult men and identified glycolysis, carbohydrate metabolism pathways as well as the pentose phosphate pathway as part of the core gut transcriptome[6]. Analysis of the transcriptomic profiles of stool samples depleted with the pan microbiome probe set confirmed confirmed the central role of these pathways in gut metabolic activity. In addition, by comparing the metatranscriptomes of infants and adults we were able to unveil additional insights on their dynamic regulation. In younger infants (<6 months of age), we saw evidence of glycolysis but also higher expression of genes involved in amino sugar metabolism (Supplemental Data File 2). The gut microbiome of these infants was largely dominated by *Bifidobacteria*, a genus well known for its ability to process human milk oligosaccharides that contains high level of N-acetylglucosamine [46]. Moreover, infant stool samples displayed higher levels of many genes involved in nucleotide metabolism, protein synthesis and cell division overall indicative of an evolving environment. In contrast, stool samples from >20 month old children showed strikingly different transcriptomic profiles and higher expression of genes involved in the catabolism of amino acids such as glycine and lysine as well as butanoate fermentation (Supplemental Data File 2). This shift paralleled a significant increase in the diversity of >20 month old children’s gut microbial profiles and acquisition of new taxa with additional metabolic capabilities (Figure 6B) that likely correlate with the introduction of solid food to the diet [47]. Amino acids are poorly absorbed in the distal colon, and hence they become an abundant and significant energy source for the gut microbiota [48, 49]. Young children (>20 months) also showed higher levels of genes associated with sporulation, likely driven by the establishment of Firmicutes such as *Clostridiales, Lachnospiraceae* and *Ruminococcus* [50] as well as motility (pili, flagella) which are found in many bacterial species and play multiple functions including promoting adhesion to the intestinal lumen (Supplemental Data File 2). Differential taxonomic expression also revealed that beyond *Bifidobacteria* younger infants had higher levels of *Escherichia, Shigella, Veillonella*, as well as >20 species of the *Enterococcus* genera. Interestingly, *Veillonella* and *Enterococcus* have been described as core components of the human milk microbiome [51, 52] while *Escherichia/Shigella* have been shown to be present at higher relative abundance in vaginally-born infants [53, 54]. Although we don’t have metadata beyond age for these infants, our findings highlight potential features of their early life environment.

Comparison of the transcriptomic profiles of infant and adult stool samples also identified numerous enzymes of the core gut transcriptome [6]. However, we noticed a striking increase in the relative expression of enzymes of the glycolysis and pentose phosphate pathways in infant samples compared to adults, suggesting a critical role for these two highly intertwined pathways in the infant gut [55]. This is in agreement with a previous study that showed the prevalence of glycolysis during the first year of life [55]. In contrast adult samples, showed increased expression of several glycosylases and sugar transporters, presumably to deal with the greater variety of carbohydrates that are part of the adult diet. Enzymes of the benzoyl-CoA pathway also showed increased expression in adult samples, which suggests that aromatic compounds are processed as carbon sources in the adult gut [56]. Interestingly, sodium benzoate is also one of the most commonly used food preservatives [57]. Additionally, adult samples expressed a number of genes involved in adhesion, biofilm formation motility, competence and quorum sensing, indicative of cells adapting and competing or cooperating to survive in an established community [58, 59].

One of the most intriguing differences we observed in metatranscriptomes was centered around stress response. Virtually all samples showed activation of stress response pathways (heat shock, cold shock, and oxidative stress), but these pathways differed significantly across age groups. Younger infants showed increased expression of many universal stress response proteins, well known for their ability to respond to a number of environmental stressors [60]. When compared to adults, infants unexpectedly showed upregulation of several cold shock proteins. We believe this difference may stem from the study design itself, as all infant stool samples were frozen and stored at −80°C prior to RNA extraction while adult samples were either extracted immediately or stabilized in DNA Genotek’s OMNIgene●GUT DNA/RNA devices. Our data highlights the importance of experimental design in metatranscriptomic analyses, since unlike DNA, gene expression profiles change quickly in response to environmental factors and stress [61].

## Conclusions

In summary, the enzymatic rRNA depletion method reported here will enable robust and accurate metatranscriptomic studies of human-associated microbial communities, allowing for detailed studies of microbiome functional activity to complement DNA-based assessments of microbial community composition.

## Declarations

### Ethics approval and consent to participate

All samples were collected under DNA Genotek’s IRB protocol (RD-PR-00087).

### Consent for publication

Not applicable.

### Availability of data and materials

Sequencing data for this project will be available through the National Center for Biotechnology Information (NCBI) Sequence Read Archive BioProject PRJNA812896.

### Author contributions

A.T, S.K., B.L.F., and D.M.G. conceived and designed experiments. A.T. designed rRNA depletion probes. A.M., J.K., D.K., F.H., V.R., E.L., and J.J. conducted experiments. A.T., S.M., J.M. analyzed data. S.K., B.L.F., and D.M.G. wrote the manuscript. All authors contributed to review and revision of the manuscript.

## Acknowledgements

We thank Andrew Cross and Shane Sontag for help with DNA sequencing and Evgueni Doukhanine for helpful comments on the manuscript. We kindly thank Dr. George Weinstock for RNA samples used to aid in the design of DPM.

## Supplemental Data Files

**Supplemental Data File 1.**

Metadata.xlsx

**Supplemental Data File 2.**

Functional_data_table_infants_vs_young_children.pdf

**Supplemental Data File 3.**

Functional_data_table_adults_vs_infants.pdf

