## Supplemental Figures for "Rational probe design for efficient rRNA depletion and improved metatranscriptomic analysis of human microbiomes"

### Supplemental Figure Legends

**Figure S1. Representative RNA quality (TapeStation) of mock RNA pool #1 as well as human stool, oral and vaginal microbiome samples used in this study.**
A) Electropherograms for total stool RNA extracted with PowerMicrobiome and RNA pool # 1 following sample analysis on an RNA screentape. B) Electropherograms for total RNA extracted from OMNIgene●ORAL (OMR-120 –tongue microbiome) and OMNIgene●VAGINAL (OMR-130) collected on high sensitivity RNA screentape.

**Figure S2. Detailed comparison of informatics tools for classifying rRNA reads from metatranscriptomes.**
Bar plots showing fractions classified as either rRNA or 'clean' (mRNA) by the various tools (bow: Bowtie 2; duk: BBDuk; sea: Seal; smr: SortMeRNA: spl: bbsplit) searched against four different databases (ar: SILVA SSU/LSU NR99; art: ar + tRNA RFAM clans; ds: SortMeRNA (4.3) default database; ss: SortMeRNA (4.3) sensitive database). The line plots represent the rRNA content of the sample. B) Read depth for the samples sequenced in this study.

**Figure S3. Detailed workflow and strategy for human microbiome rRNA depletion probe design.**
A) Detailed flowchart of the iterative probe design process used to generate HMv1 and HMv2 custom rRNA depletion probe sets. The steps highlighted in blue are described in more detail in B) and C). B) Following depletion of donor samples with DP1 and alignment of fastq files to the SilvaDB, regions of rRNAs that have over 500x coverage are collected. A representative donor sample is visualized in IGV (upper panel) showing both undepleted and DP1 depleted sequencing reads with alignment of the DP1 probes across the bottom of the panel. Regions where DP1 probes do not align corresponds to high rRNA read coverage (dotted lines). The rRNA regions with >500x coverage were collected and their length was plotted vs frequency observed (two examples shown in lower panels). Most of these regions are <200 nt in length. C) A heatmap showing the top 50 most abundant regions from each donor that were pooled and pairwise alignment was performed. Red indicated regions with high percent alignment and blue represents low percent alignment. Gray represent zero percent alignment. If two regions aligned >80% only one region was chosen to reduce redundancy. Therefore, red areas of the heatmap resulted in fewer probes designed, while blue or gray areas required more probes to be generated.

**Figure S4. Functional profiles of adult fecal metatranscriptomes.**
A) RefSeq Functions. B) SEED Functions.

**Figure S5. Comparing the rRNA depletion efficiency of the original Ribo-Zero Gold (Epidemiology) kits or Ribo-Zero Plus with DP1 or HMv1 probe sets from ATCC mock community samples.**
A) Depletion from ATCC skin samples is more efficient when using the Ribo-Zero Gold (Epidemiology) kit (RZBact) than Ribo-Zero Plus (RZ + DP1). However, substituting DP1 with the HMv1 probe set results in efficient rRNA depletion (HMv1). B) Depletion of ATCC gut samples is effective using only DP1 and demonstrates that Ribo-Zero Plus has better reproducibility than the traditional hybrid capture method which is prone to contamination caused by inadvertent carry over of depletion beads during the protocol.

**Figure S6. RT-qPCR based assessment of vaginal microbiome *Lactobacillus* content and additional oral rRNA depletion data.**

A) RT-qPCR quantification of *Lactobacillus*-dominated and non-*Lactobacillus*-dominated samples and controls. B) rRNA depletion of additional tongue microbiome samples using the HMv1 probes.

**Figure S7. ERCC spike-in depletion and mapping data.**
A) rRNA depletion for indicated ERCC spike-in samples. B) Proportion of reads aligning to all ERCC spike-ins in indicated samples. C) For the reads aligning to the ERCC spike-ins, the proportion of reads aligning to each ERCC spike-in for indicated samples.

**Figure S8. rRNA depletion efficiency in stool samples of donors between 4 and 33 months old.**
A) Percentage of reads mapping to eukaryotic and prokaryotic rRNAs vs coding sequences in infant and child stool samples depleted with Ribo-Zero Plus with HMv1 probes. Replicate rRNA depletion reactions were performed for each sample. B) Percentage of reads mapping to eukaryotic and prokaryotic rRNAs vs. other sequences in four infant stool samples depleted with Ribo-Zero Plus with HMv1 or HMv2 compared to undepleted samples. Replicate rRNA depletion reactions were performed for each condition. Boxed bar plot on top of panel B represents remaining bacterial rRNA (LSU & SSU) with a dashed line at the 50% mark.

**Figure S9. Comparison of rRNA depletion efficiency of adult stool RNA samples using the HMv1 and HMv2 custom probe sets**
Percentage of reads mapping to eukaryotic and prokaryotic rRNAs vs. other sequences in adult stool samples depleted with Ribo-Zero Plus with HMv1 or HMv2. the same 10 healthy adult samples were used as in Figure 3.

**Figure S10. Comparison of number of features detected for infant samples depleted with different methods.**
Number of reads, or taxonomic or functional features detected for undepleted, HMv1-depleted, or HMv2-depleted infant samples.

**Figure S11. Infant versus adult metatranscriptomic profiles.**
A) Adult and infant RefSeq taxa. B) Adult and infant CAZyme profiles.

**Figure S12. Infant versus adult metatranscriptomic profiles.**
Effects plot showing functional categories that exhibit differential expression between adult and infant samples.

**Figure S13. Volcano plots of adult and infant taxa or function.**
Taxa (A, C) or functional groups (B, D) that are significantly different in the indicated comparison are colored in red, non-significant differences are in black. A) Infant taxa from samples <6 months vs. >20 months, B) Infant RefSeq functions from samples <6 months vs. >20 months, C) Infant vs. adult taxa, D) Infant vs. adult RefSeq functions.

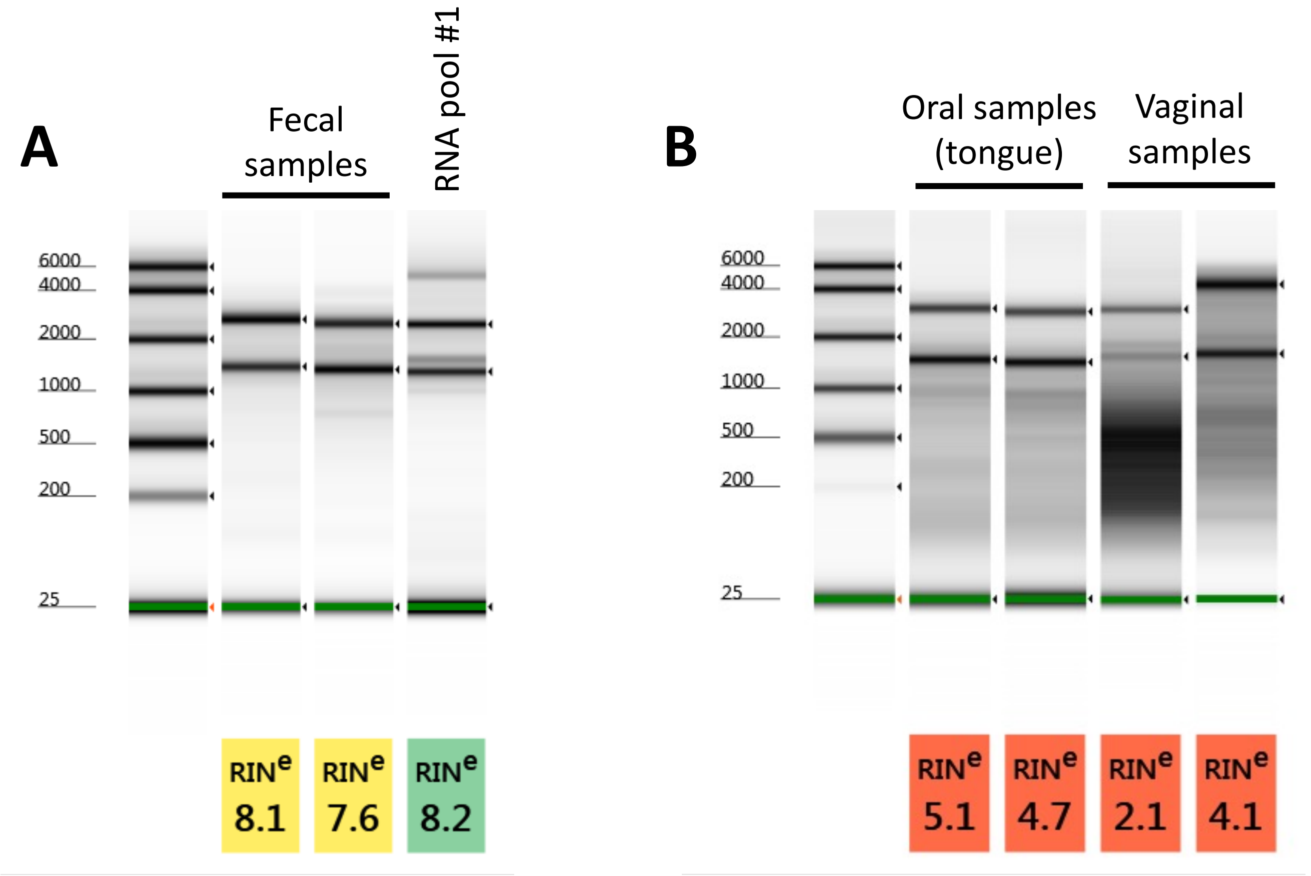
A
Fecal samples
RNA pool #1
6000
4000
2000
1000
500
200
25
RIN^e 8.1
RIN^e 7.6
RIN^e 8.2
B
Oral samples (tongue)
Vaginal samples
6000
4000
2000
1000
500
200
25
RIN^e 5.1
RIN^e 4.7
RIN^e 2.1
RIN^e 4.1

Figure S1

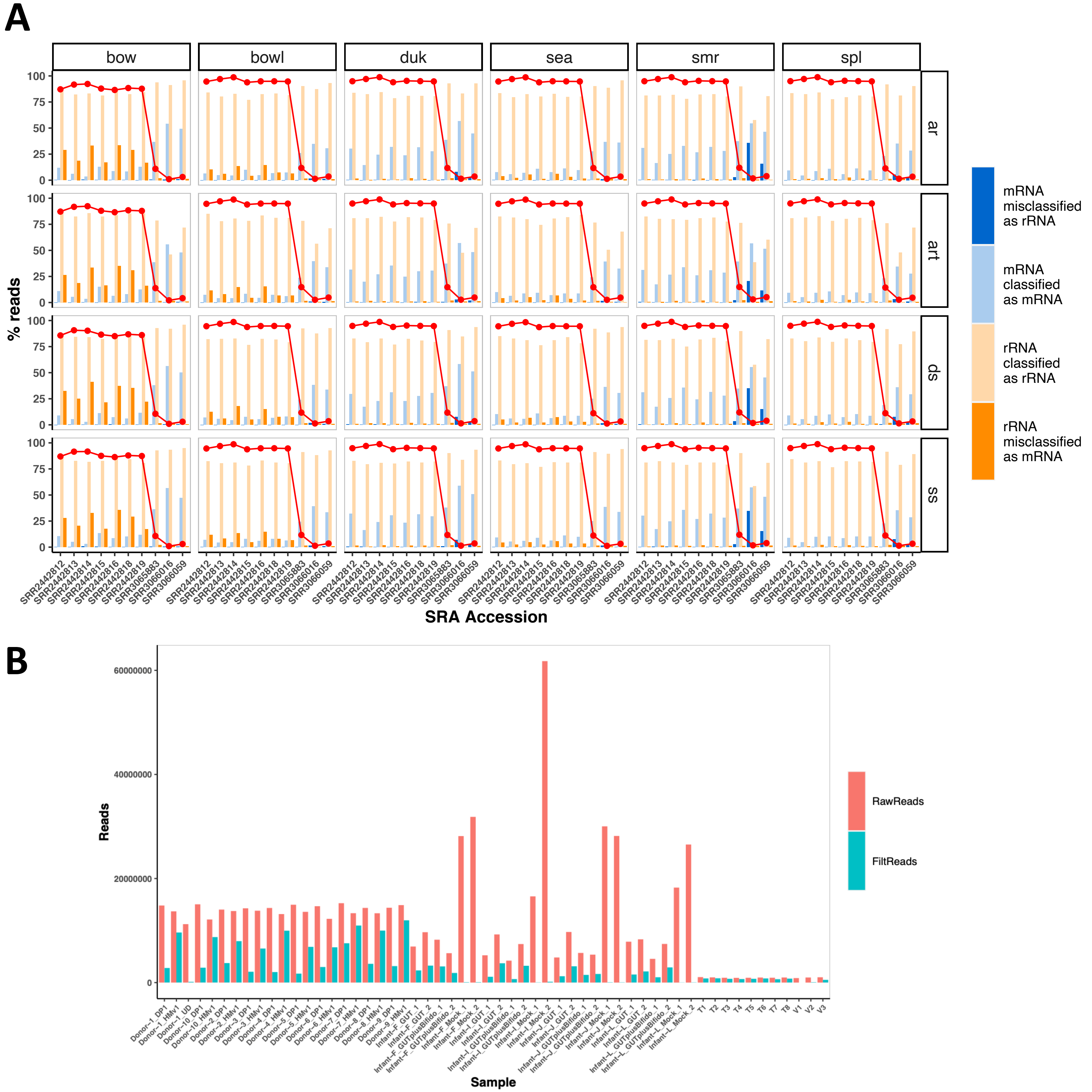
A
bow
bowl
duk
sea
smr
spl
ar
art
ds
ss
% reads
100
75
50
25
0
SRR2442812
SRR2442813
SRR2442814
SRR2442815
SRR2442816
SRR2442818
SRR2442819
SRR3065883
SRR3066016
SRR3066059
SRA Accession
mRNA misclassified as rRNA
mRNA classified as mRNA
rRNA classified as rRNA
rRNA misclassified as mRNA
B
Reads
60000000
40000000
20000000
0
Sample
RawReads
FiltReads

Figure S2

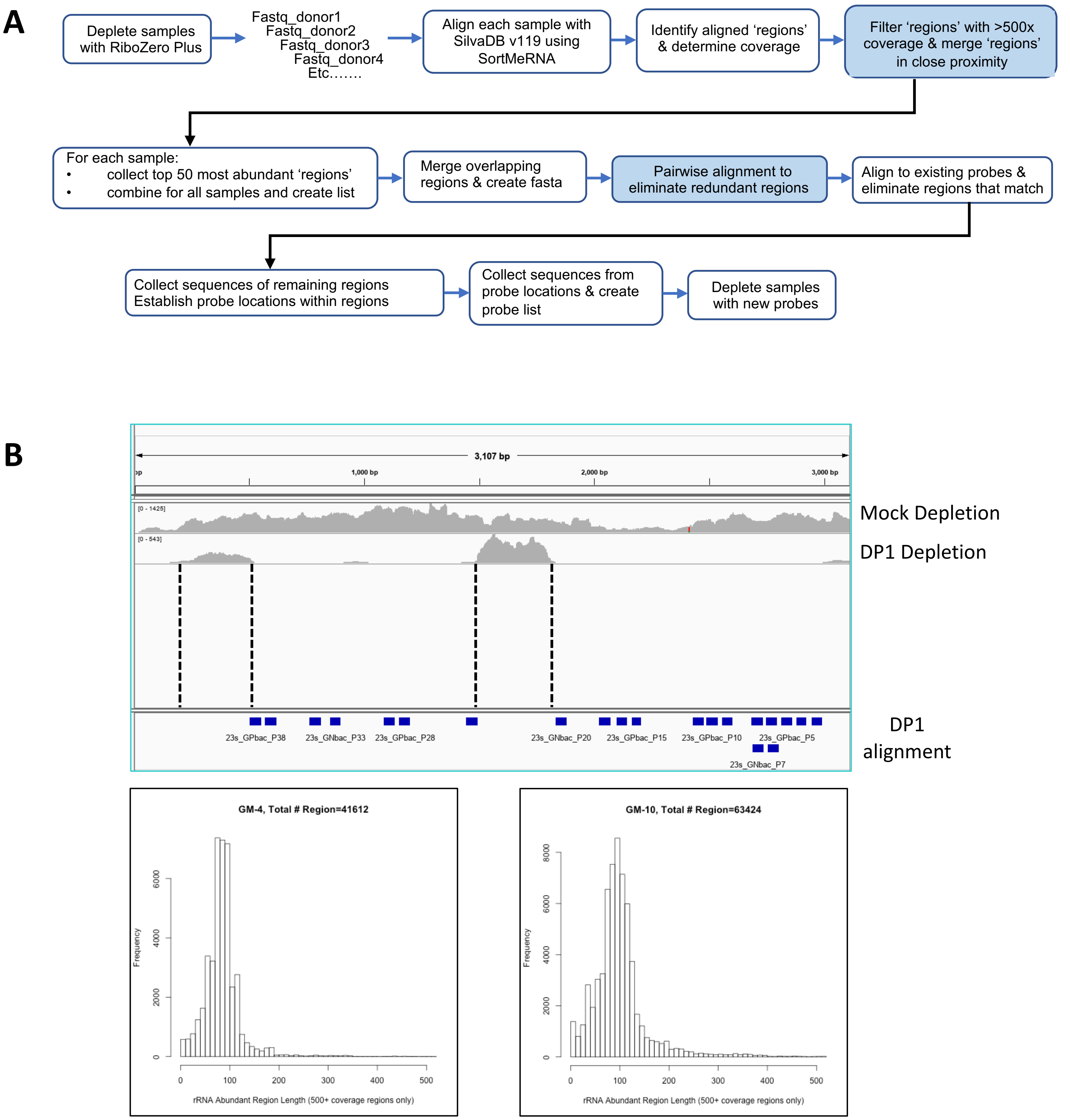
A
Deplete samples with RiboZero Plus
Fastq_donor1
Fastq_donor2
Fastq_donor3
Fastq_donor4
Etc…….
Align each sample with SilvaDB v119 using SortMeRNA
Identify aligned ‘regions’ & determine coverage
Filter ‘regions’ with >500x coverage & merge ‘regions’ in close proximity
For each sample:
• collect top 50 most abundant ‘regions’
• combine for all samples and create list
Merge overlapping regions & create fasta
Pairwise alignment to eliminate redundant regions
Align to existing probes & eliminate regions that match
Collect sequences of remaining regions
Establish probe locations within regions
Collect sequences from probe locations & create probe list
Deplete samples with new probes
B
3,107 bp
1,000 bp
2,000 bp
3,000 bp
[0 - 1425]
[0 - 543]
Mock Depletion
DP1 Depletion
23s_GPbac_P38
23s_GNbac_P33
23s_GPbac_P28
23s_GNbac_P20
23s_GPbac_P15
23s_GPbac_P10
23s_GPbac_P5
23s_GNbac_P7
DP1 alignment
GM-4, Total # Region=41612
Frequency
0
2000
4000
6000
0
100
200
300
400
500
rRNA Abundant Region Length (500+ coverage regions only)
GM-10, Total # Region=63424
Frequency
0
2000
4000
6000
8000
0
100
200
300
400
500
rRNA Abundant Region Length (500+ coverage regions only)

Figure S3

C

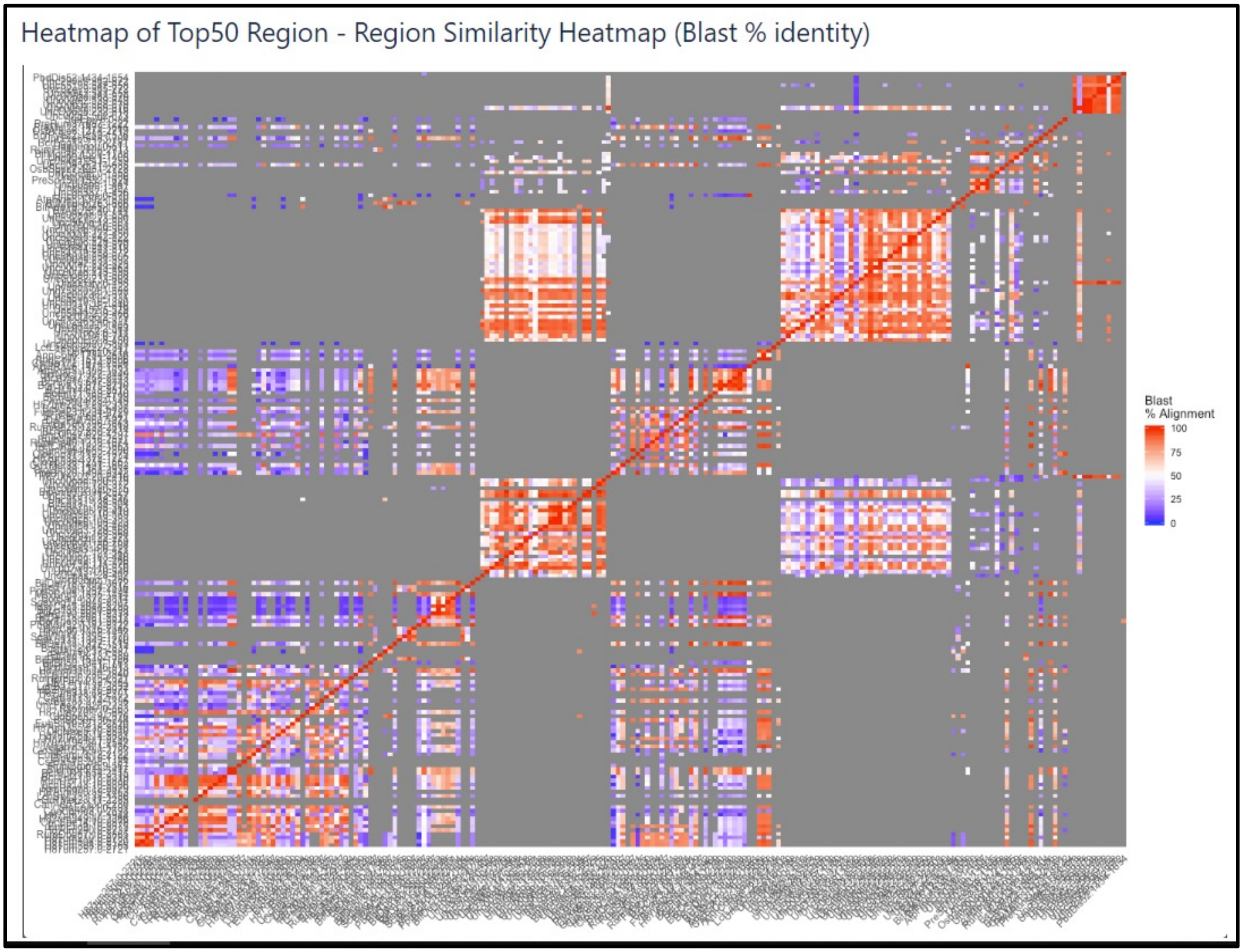

Heatmap of Top50 Region - Region Similarity Heatmap (Blast % identity)
Blast
% Alignment
100
75
50
25
0

A

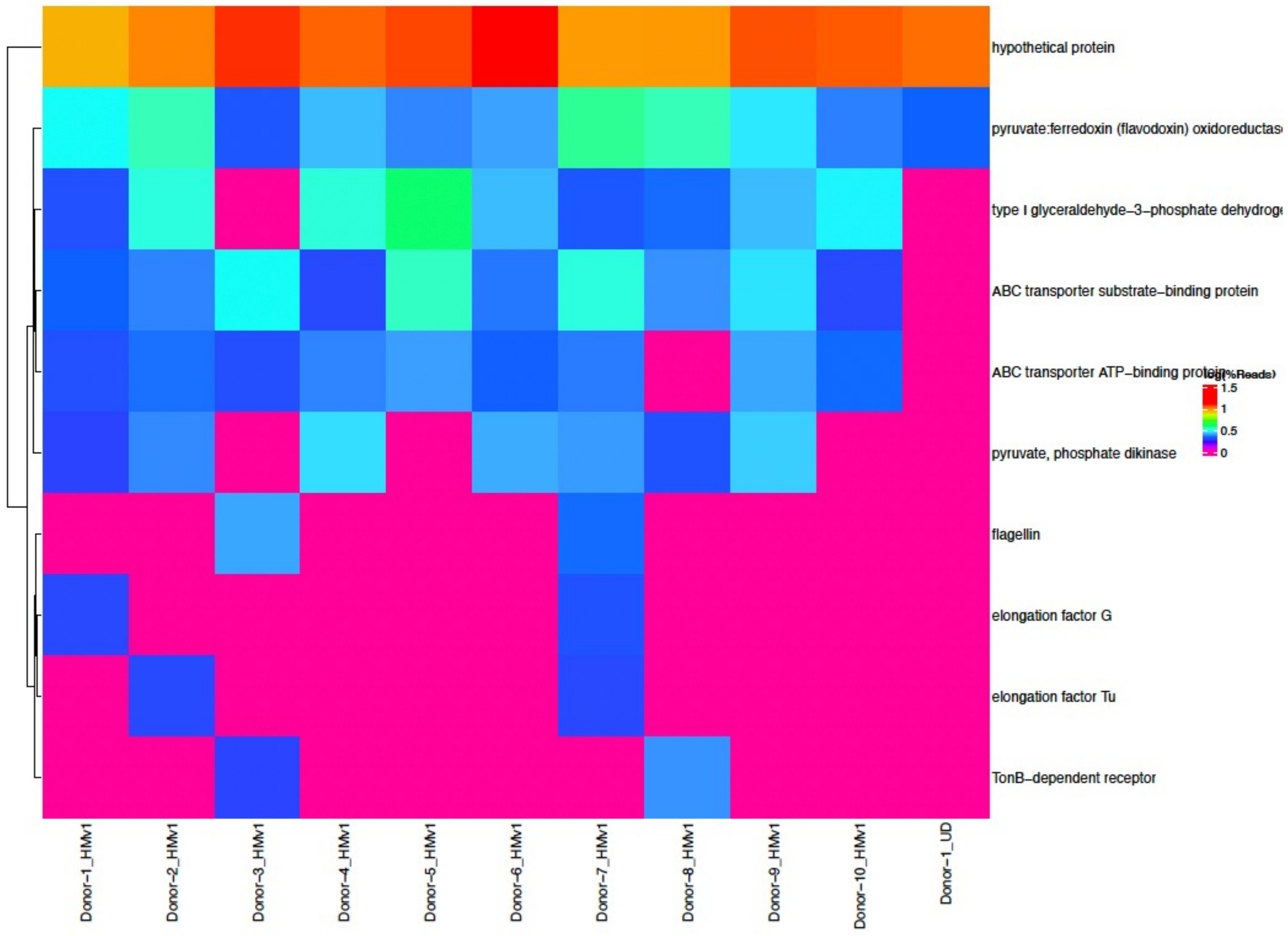
hypothetical protein
pyruvate:ferredoxin (flavodoxin) oxidoreductase
type I glyceraldehyde-3-phosphate dehydrog
ABC transporter substrate-binding protein
ABC transporter ATP-binding protein
log(%Reads)
1.5
1
0.5
0
pyruvate, phosphate dikinase
flagellin
elongation factor G
elongation factor Tu
TonB-dependent receptor
Donor-1_HMv1
Donor-2_HMv1
Donor-3_HMv1
Donor-4_HMv1
Donor-5_HMv1
Donor-6_HMv1
Donor-7_HMv1
Donor-8_HMv1
Donor-9_HMv1
Donor-10_HMv1
Donor-1_UD

B

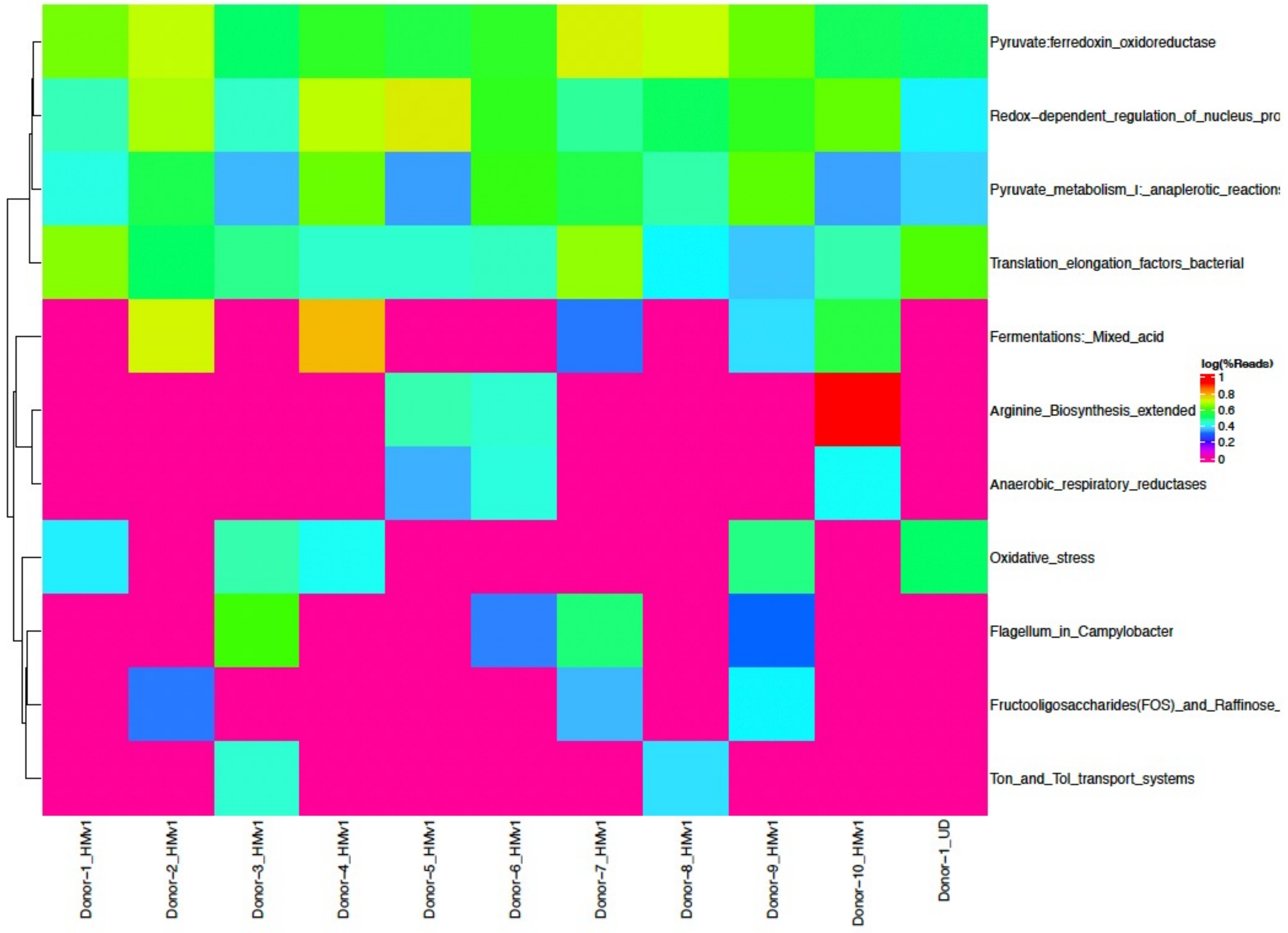
Pyruvate:ferredoxin_oxidoreductase
Redox-dependent_regulation_of_nucleus_pro
Pyruvate_metabolism_I:_anaplerotic_reaction
Translation_elongation_factors_bacterial
Fermentations:_Mixed_acid
log(%Reads)
1
0.8
0.6
0.4
0.2
0
Arginine_Biosynthesis_extended
Anaerobic_respiratory_reductases
Oxidative_stress
Flagellum_in_Campylobacter
Fructooligosaccharides(FOS)_and_Raffinose_
Ton_and_Tol_transport_systems
Donor-1_HMv1
Donor-2_HMv1
Donor-3_HMv1
Donor-4_HMv1
Donor-5_HMv1
Donor-6_HMv1
Donor-7_HMv1
Donor-8_HMv1
Donor-9_HMv1
Donor-10_HMv1
Donor-1_UD

Figure S4

**A.** ATCC skin

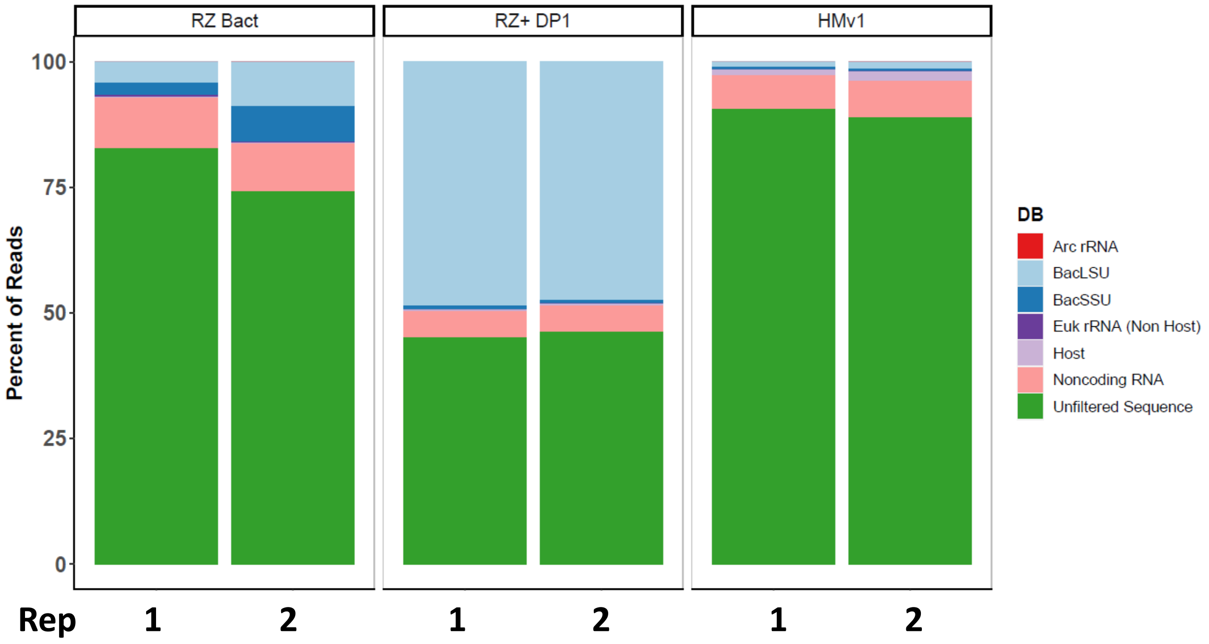


**B.** ATCC gut

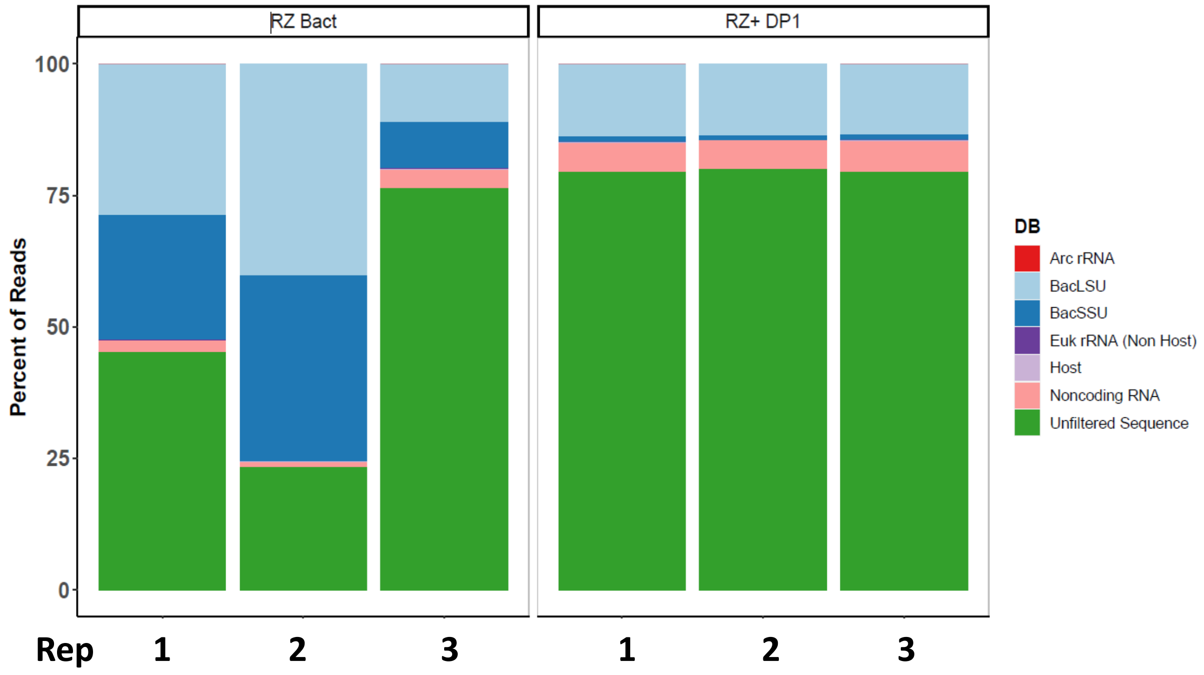


Figure S5

A

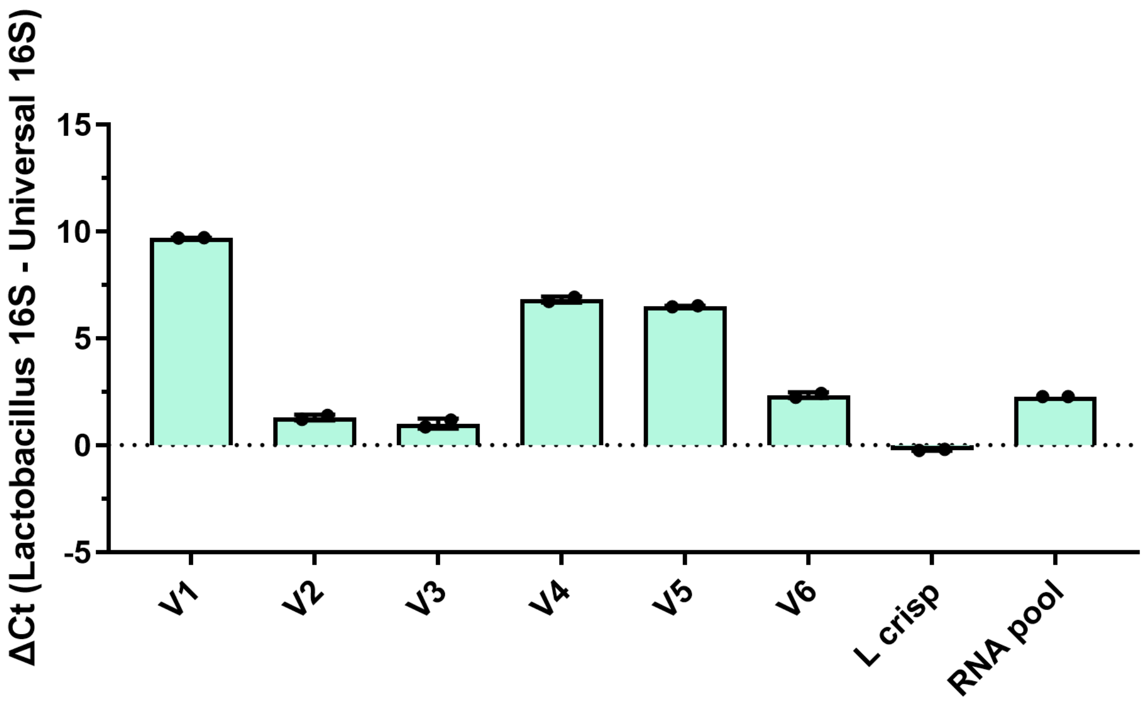
ΔCt (Lactobacillus 16S - Universal 16S)
15
10
5
0
-5
V1
V2
V3
V4
V5
V6
L crisp
RNA pool

B

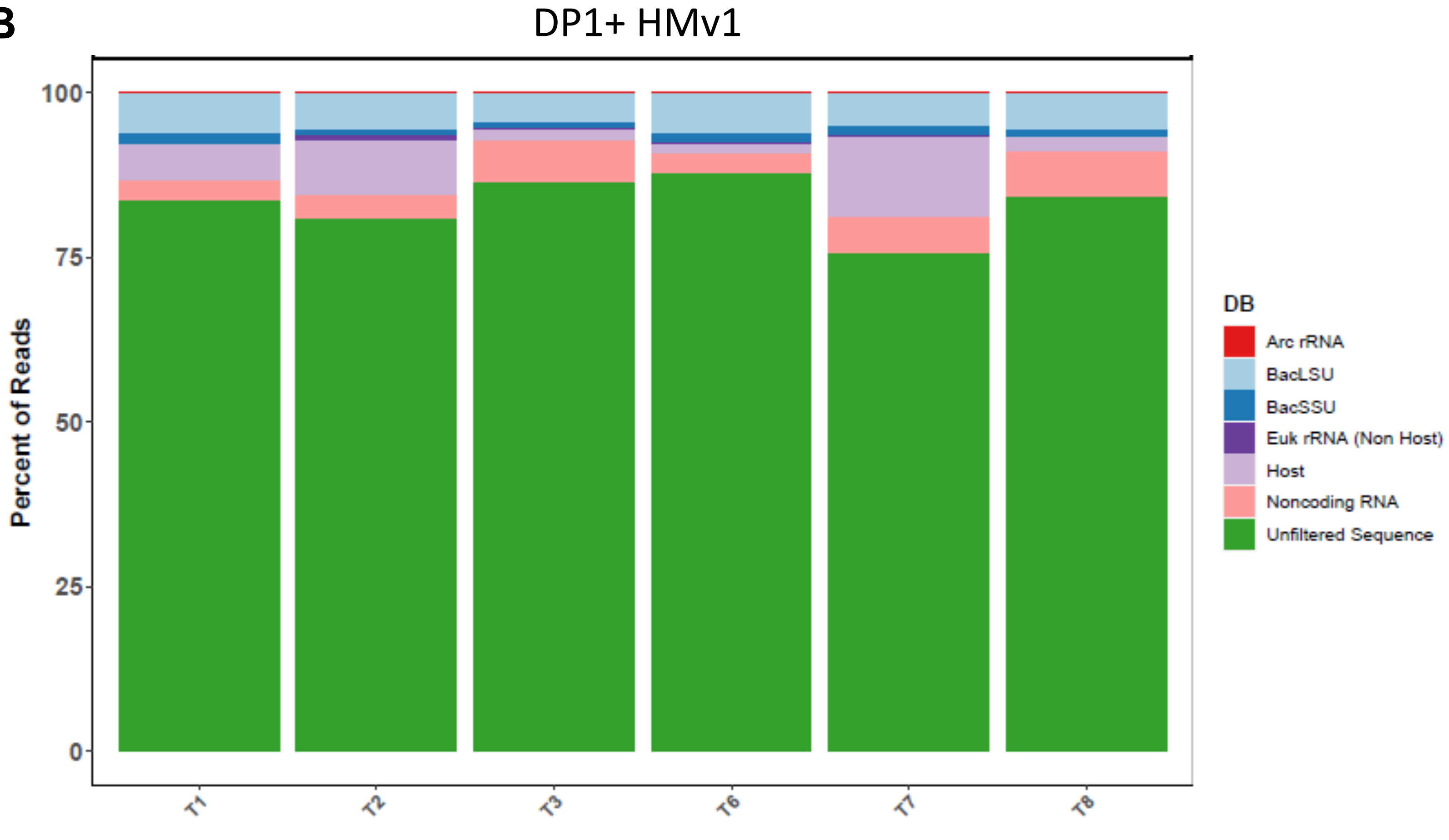
DP1+ HMv1
Percent of Reads
100
75
50
25
0
T1
T2
T3
T6
T7
T8
DB
Arc rRNA
BacLSU
BacSSU
Euk rRNA (Non Host)
Host
Noncoding RNA
Unfiltered Sequence

Figure S6

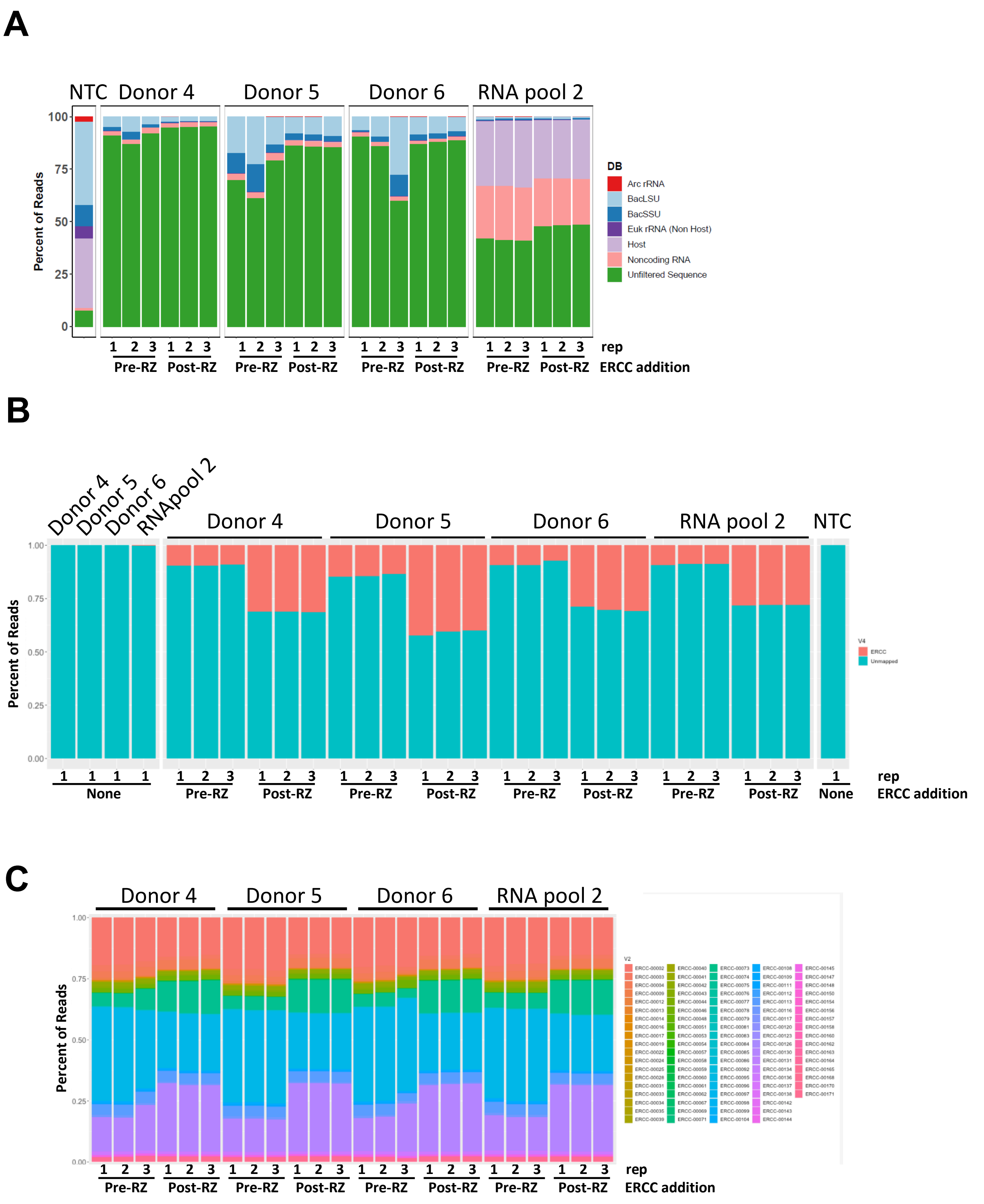
A
NTC
Donor 4
Donor 5
Donor 6
RNA pool 2
Percent of Reads
100
75
50
25
0
DB
Arc rRNA
BacLSU
BacSSU
Euk rRNA (Non Host)
Host
Noncoding RNA
Unfiltered Sequence
1 2 3 1 2 3
Pre-RZ Post-RZ
rep
ERCC addition
B
Donor 4
Donor 5
Donor 6
RNApool 2
RNA pool 2
NTC
Percent of Reads
1.00
0.75
0.50
0.25
0.00
V4
ERCC
Unmapped
None
Pre-RZ
Post-RZ
rep
ERCC addition
C
Donor 4
Donor 5
Donor 6
RNA pool 2
Percent of Reads
1.00
0.75
0.50
0.25
0.00
Pre-RZ
Post-RZ
rep
ERCC addition

Figure S7

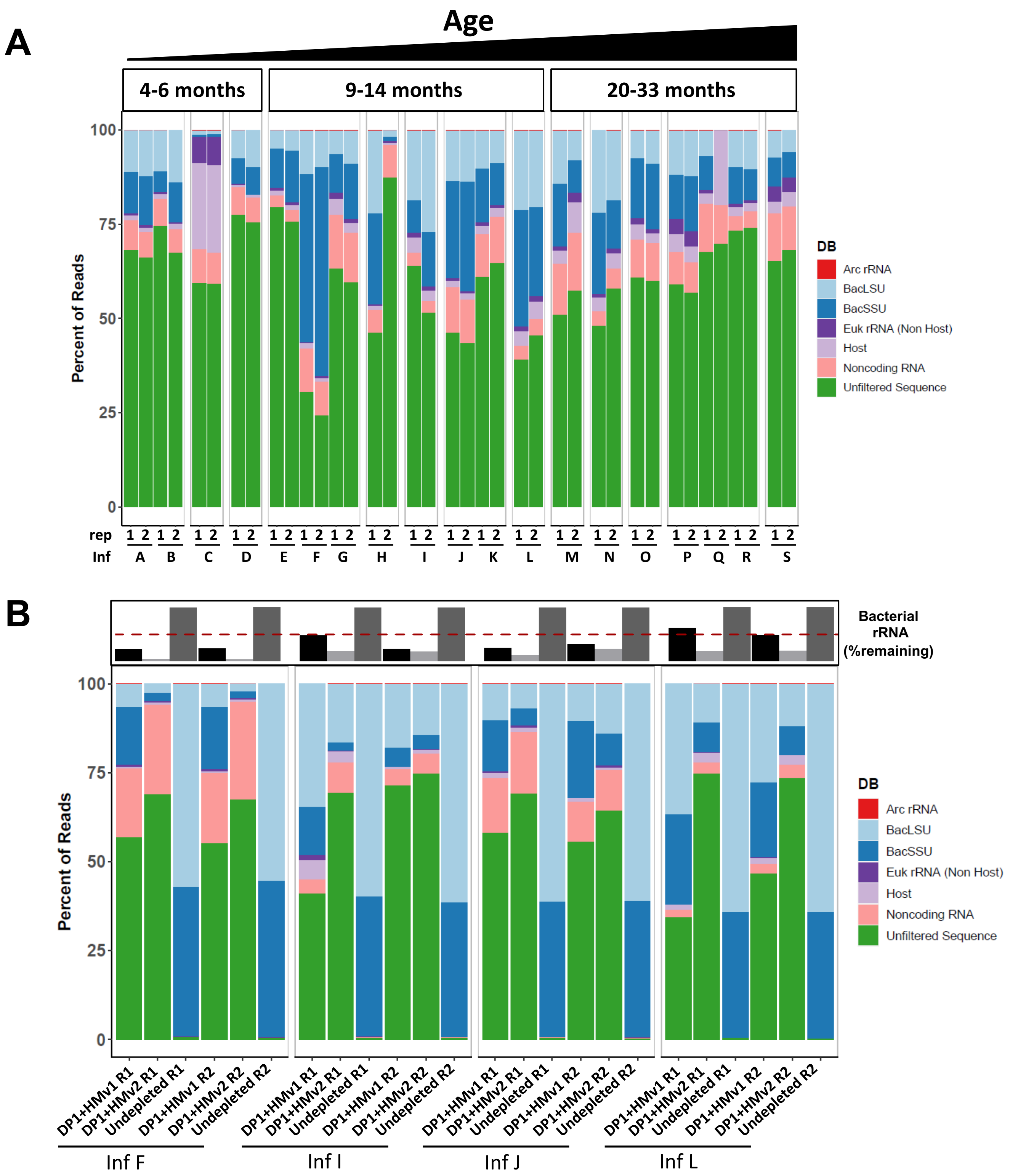

A
Age
4-6 months
9-14 months
20-33 months
Percent of Reads
100
75
50
25
0
rep
Inf
A B C D E F G H I J K L M N O P Q R S
DB
Arc rRNA
BacLSU
BacSSU
Euk rRNA (Non Host)
Host
Noncoding RNA
Unfiltered Sequence
B
Bacterial rRNA (%remaining)
DP1+HMv1 R1
DP1+HMv2 R1
Undepleted R1
DP1+HMv1 R2
DP1+HMv2 R2
Undepleted R2
Inf F
Inf I
Inf J
Inf L


Figure S8

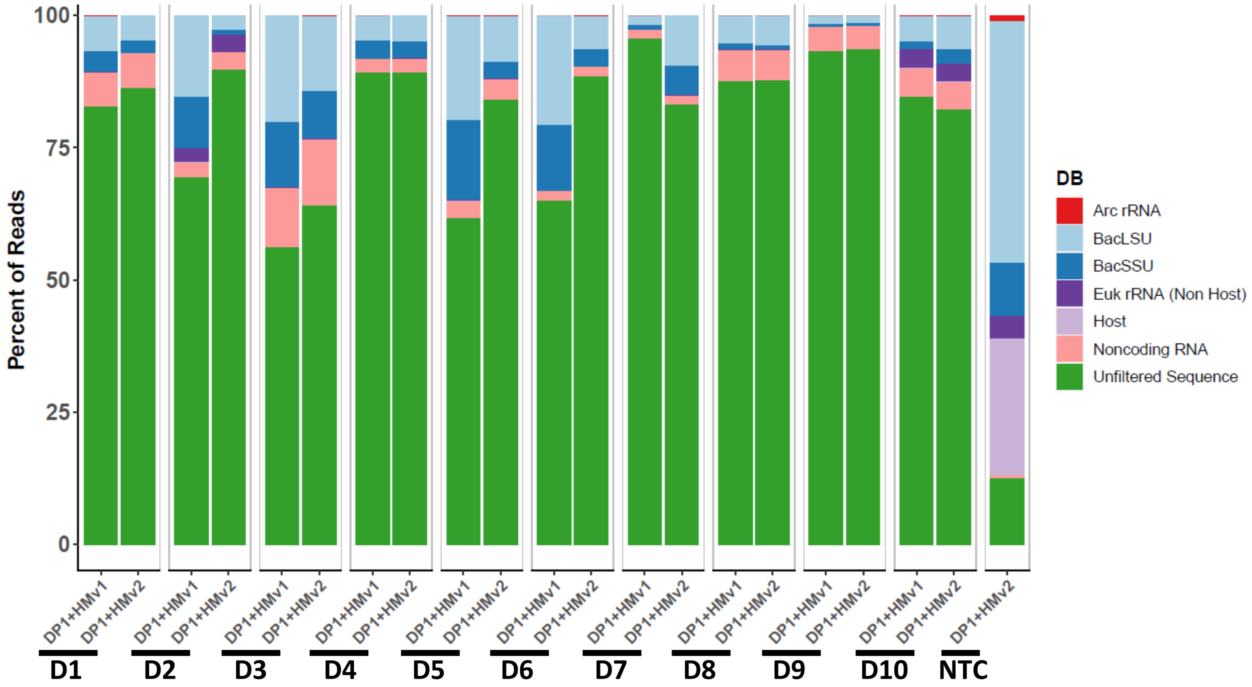
Percent of Reads
100
75
50
25
0
DP1+HMv1
DP1+HMv2
D1
D2
D3
D4
D5
D6
D7
D8
D9
D10
NTC
DB
Arc rRNA
BacLSU
BacSSU
Euk rRNA (Non Host)
Host
Noncoding RNA
Unfiltered Sequence

Figure S9

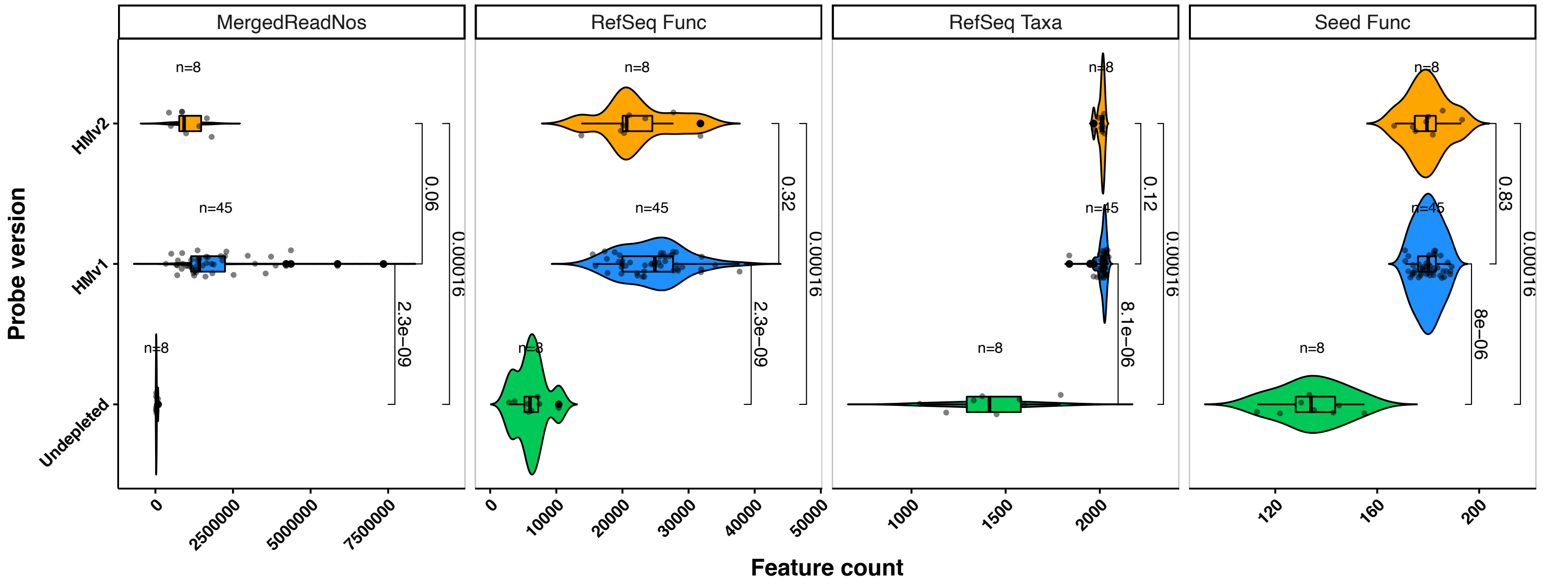

MergedReadNos
RefSeq Func
RefSeq Taxa
Seed Func
Probe version
HMv2
HMv1
Undepleted
n=8
n=45
n=8
0.06
2.3e−09
0.00016
0.32
0.12
8.1e−06
0.83
8e−06
0
2500000
5000000
7500000
10000
20000
30000
40000
50000
1000
1500
2000
120
160
200
Feature count


Figure S10

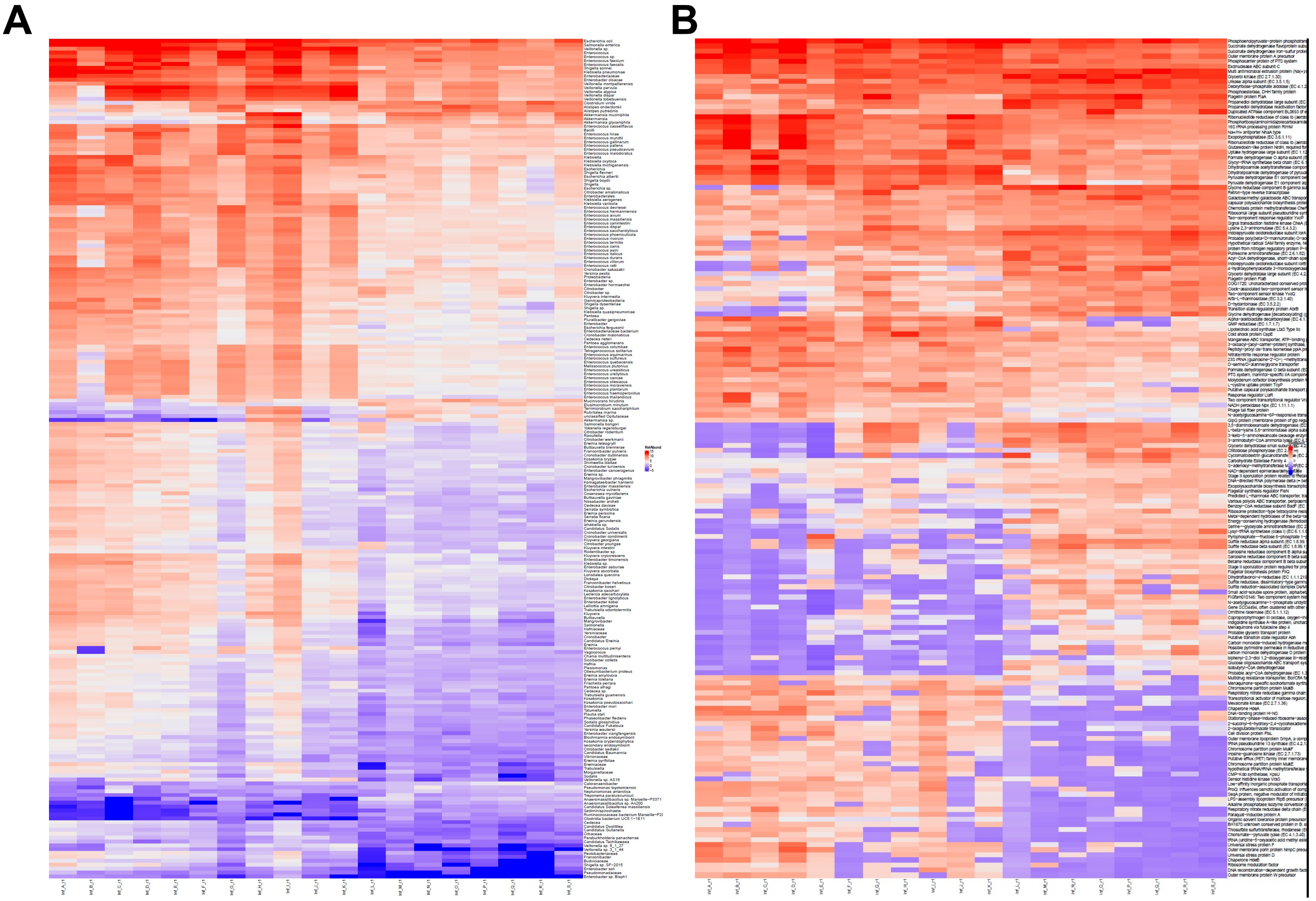

Figure S11

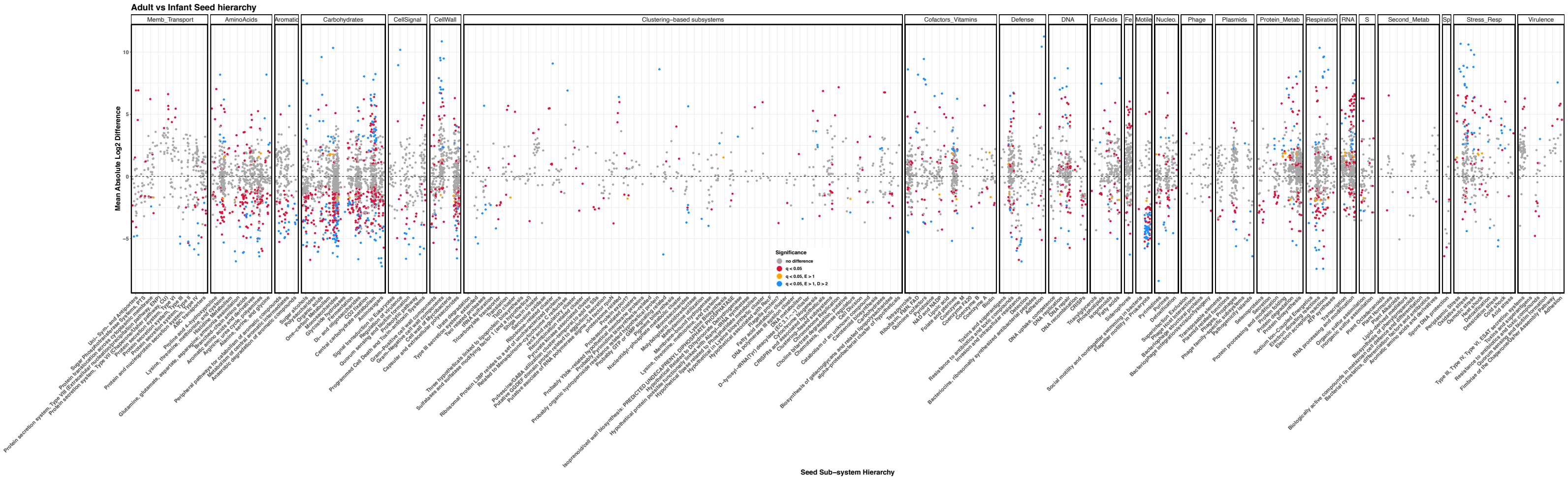
Adult vs Infant Seed hierarchy
Memb_Transport
AminoAcids
Aromatic
Carbohydrates
CellSignal
CellWall
Clustering−based subsystems
Cofactors_Vitamins
Defense
DNA
FatAcids
Fe
Motile
Nucleo.
Phage
Plasmids
Protein_Metab
Respiration
RNA
S
Second_Metab
Sp
Stress_Resp
Virulence
Mean Absolute Log2 Difference
10
5
-5
Significance
no difference
q < 0.05
q < 0.05, E > 1
q < 0.05, E > 1, D > 2
Seed Sub−system Hierarchy

Figure S12

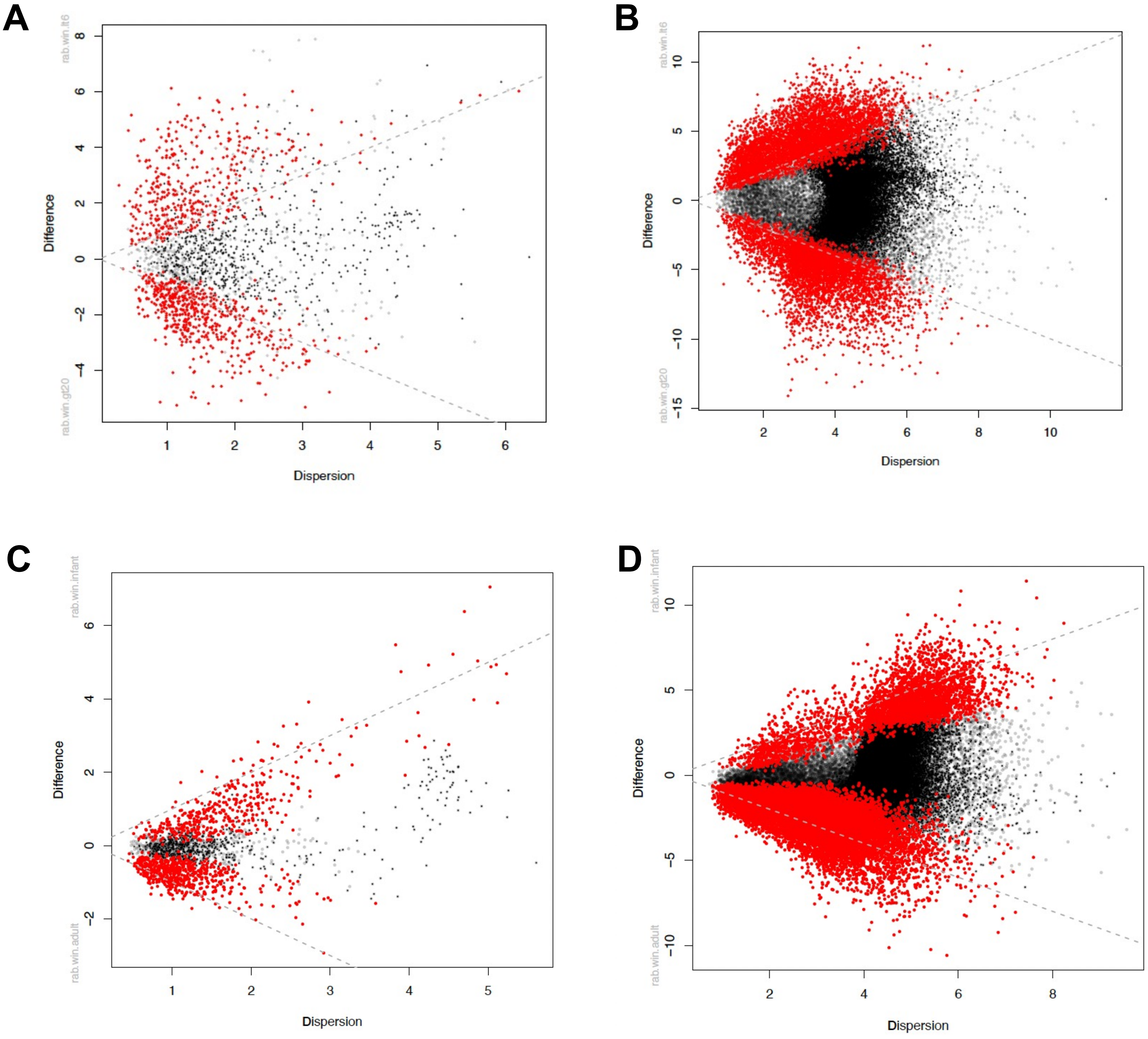
A
rab.win.lt6
rab.win.gt20
Difference
Dispersion
B
rab.win.lt6
rab.win.gt20
Difference
Dispersion
C
rab.win.infant
rab.win.adult
Difference
Dispersion
D
rab.win.infant
rab.win.adult
Difference
Dispersion

Figure S13
