## Supplemental File 2 for "Rational probe design for efficient rRNA depletion and improved metatranscriptomic analysis of human microbiomes"

| | | Effect size |
|---|---|---|
| Carbohydrate metabolism | UDP-N-acetylglucosamine 4-epimerase (EC 5.1.3.7) | 3.28 |
| | N-acetylglucosamine-6P-responsive transcriptional repressor NagC, ROK family | 2.74 |
| | Beta N-acetyl-glucosaminidase (EC 3.2.1.52) | 2.67 |
| | Beta-hexosaminidase (EC 3.2.1.52) | 2.52 |
| | Galactokinase (EC 2.7.1.6) | 2.19 |
| | Phosphoglycerate mutase (EC 5.4.2.1) | 2.05 |
| | Aldose 1-epimerase family protein YeaD | 1.95 |
| | Glucosamine-6-phosphate deaminase (EC 3.5.99.6) | 1.88 |
| | 6-phosphofructokinase class II (EC 2.7.1.11) | 1.87 |
| | Sialidase (EC 3.2.1.18) | 1.69 |
| | Glucokinase (EC 2.7.1.2) | 1.66 |
| | Fructose-1,6-bisphosphatase, type I (EC 3.1.3.11) | 1.55 |
| | Pyrophosphate--fructose 6-phosphate 1-phosphotransferase, alpha subunit (EC 2.7.1.90) | -1.97 |
| | 2-dehydro-3-deoxygluconate kinase (EC 2.7.1.45) | -2.04 |
| | Phosphoenolpyruvate carboxykinase [GTP] (EC 4.1.1.32) | -2.59 |
| | Fructose-bisphosphate aldolase, archaeal class I (EC 4.1.2.13) | -2.73 |
| | Butyryl-CoA dehydrogenase (EC 1.3.99.2) | -3.62 |
| | Methylglyoxal synthase (EC 4.2.3.3) | -4.61 |
| Amino acid metabolism | Serine--glyoxylate aminotransferase (EC 2.6.1.45) | -2.40 |
| | 3-ketoacyl-CoA thiolase [isoleucine degradation] (EC 2.3.1.16) | -2.42 |
| | Sarcosine reductase component B beta subunit (EC 1.21.4.3) | -2.47 |
| | N-methylhydantoinase A (EC 3.5.2.14) | -2.51 |
| | Sarcosine reductase component B alpha subunit (EC 1.21.4.3) | -2.58 |
| | Glycine reductase component B alpha subunit (EC 1.21.4.2) | -2.66 |
| | Lysine 2,3-aminomutase (EC 5.4.3.2) | -3.31 |
| | Butyrate-acetoacetate CoA-transferase subunit B (EC 2.8.3.9) | -3.31 |
| | Glycine/sarcosine/betaine reductase protein A | -3.36 |
| | L-beta-lysine 5,6-aminomutase beta subunit (EC 5.4.3.3) | -3.65 |
| | 3,5-diaminobexanoate dehydrogenase (EC 1.4.1.11) | -3.95 |

| | | |
|---|---|---|
| | 3-aminobutyrl-CoA ammonia lyase (EC 4.3.1.14) | -4.24 |
| | L-beta-lysine 5,6-aminomutase alpha subunit (EC 5.4.3.3) | -4.33 |
| | 3-keto-5-aminohexanoate cleavage enzyme | -4.34 |
| Motility & sporulation | Type IV fimbrial assembly protein PilC | -2.00 |
| | Stage V sporulation protein AC (SpoVAC) | -2.05 |
| | Flagellar biosynthesis protein FliQ | -2.07 |
| | Stage III sporulation protein D | -2.08 |
| | Flagellin protein FlaB | -2.14 |
| | Dipicolinate synthase subunit B | -2.22 |
| | Transition state regulatory protein AbrB | -2.59 |
| | Stage V sporulation protein T, AbrB family transcriptional regulator (SpoVT) | -2.63 |
| | Type IV pilin PilA | -2.69 |
| | Stage II sporulation protein required for processing of pro-sigma-E (SpoIIR) | -3.01 |
| Stress response | Cold shock protein CspE | 4.01 |
| | Chaperone HdeB | 3.38 |
| | Chaperone HdeA | 2.79 |
| | Universal stress protein F | 2.50 |
| | Error-prone repair protein UmuD | 2.30 |
| | Alkyl hydroperoxide reductase protein C (EC 1.6.4.-) | 2.24 |
| | Chaperone-modulator protein CbpM | 2.11 |
| | Universal stress protein D | 2.07 |
| | Universal stress protein A | 2.03 |
| | Paraquat-inducible protein A | 1.95 |
| | Universal stress protein G | 1.88 |
| | Universal stress protein B | 1.75 |
| | Two-component response regulator YvcP | -2.31 |
| | Two-component sensor kinase YvcQ | -2.34 |
| | Two-component sensor histidine kinase BceS | -2.36 |
| | anti sigma b factor antagonist RsbV | -2.36 |
| | anti-sigma B factor RsbT | -3.34 |
| [illegible] pyruvate metabolism | Succinyl-CoA ligase [ADP-forming] beta chain (EC 6.2.1.5) | 2.63 |
| | Succinate dehydrogenase flavoprotein subunit (EC 1.3.99.1) | 2.42 |
| | Succinyl-CoA ligase [ADP-forming] alpha chain (EC 6.2.1.5) | 2.41 |
| | Succinate dehydrogenase iron-sulfur protein (EC 1.3.99.1) | 2.29 |
| | Dihydrolipoamide dehydrogenase (EC 1.8.1.4) | 2.07 |
| | Dihydrolipoamide dehydrogenase of 2-oxoglutarate dehydrogenase (EC 1.8.1.4) | 1.96 |

| | | |
|---|---|---|
| Succinate | Dihydrolipoamide acetyltransferase component of pyruvate dehydrogenase complex (EC 2.3.1.12) | 1.77 |
| | Pyruvate dehydrogenase E1 component alpha subunit (EC 1.2.4.1) | 1.69 |
| | Pyruvate dehydrogenase E1 component beta subunit (EC 1.2.4.1) | 1.64 |
