## Supplemental File 3 for "Rational probe design for efficient rRNA depletion and improved metatranscriptomic analysis of human microbiomes"

| | | Effect size |
|---|---|---|
| Carbohydrate metabolism | Mannonate dehydratase (EC 4.2.1.8) | -1.47 |
| | Gluconate dehydratase (EC 4.2.1.39) | -1.47 |
| | Transaldolase (EC 2.2.1.2) | 1.36 |
| | Ribose 5-phosphate isomerase A (EC 5.3.1.6) | 1.45 |
| | Glucose dehydrogenase, PQQ-dependent (EC 1.1.5.2) | 1.57 |
| | Aerobic glycerol-3-phosphate dehydrogenase (EC 1.1.5.3) | 1.74 |
| | Enolase (EC 4.2.1.11) | 1.94 |
| | Phosphoglycerate mutase (EC 5.4.2.1) | 1.96 |
| | Glucose-6-phosphate 1-dehydrogenase (EC 1.1.1.49) | 2.00 |
| | 6-phosphogluconate dehydrogenase, decarboxylating (EC 1.1.1.44) | 2.06 |
| Pyruvate and benzoate metabolism | Benzoyl-CoA reductase subunit BadG (EC 1.3.99.15) | -1.73 |
| | Benzoyl-CoA reductase subunit BadF (EC 1.3.99.15) | -1.59 |
| | Benzoyl-CoA reductase subunit BadE (EC 1.3.99.15) | -1.45 |
| | Benzoyl-CoA reductase subunit BadD (EC 1.3.99.15) | -1.33 |
| | Carbon monoxide dehydrogenase large chain (EC 1.2.99.2) without typical motifs | -1.30 |
| | 3-oxoadipate CoA-transferase subunit A (EC 2.8.3.6) | -1.29 |
| | Homocitrate synthase (EC 2.3.3.14) | -1.28 |
| | Dihydrolipoamide acetyltransferase component of pyruvate dehydrogenase complex (EC 2.3.1.12) | 1.39 |
| | Pyruvate oxidase [ubiquinone, cytochrome] (EC 1.2.2.2) | 1.50 |
| | 2-methylisocitrate dehydratase (EC 4.2.1.99) | 1.50 |
| | Pyruvate dehydrogenase E1 component (EC 1.2.4.1) | 1.72 |
| te membrane transport | Melibiose carrier protein, Na+/melibiose symporter | -1.61 |
| | Unknown carbohydrate transporter from TRAP family, substrate-binding component UctP | -1.61 |
| | Unspecified monosaccharide ABC transport system, permease component 2 | -1.57 |
| | Beta-glucoside ABC transport system, sugar-binding protein | -1.56 |
| | Predicted glycosylase TM1225 | -1.44 |
| | Unknown carbohydrate transporter from TRAP family, large transmembrane component UctQ | -1.43 |

negative effect size va
positive effect size val

| | | |
|---|---|---|
| Glycosylases and carbohydra[illegible] | COG2152 predicted glycoside hydrolase | -1.43 |
| | Predicted glycosylase, COG2152 | -1.41 |
| | Various polyols ABC transporter, periplasmic substrate-binding protein | -1.40 |
| | Unspecified monosaccharide ABC transport system, substrate-binding component | -1.39 |
| | Unspecified monosaccharide ABC transport system, permease component Ib (FIG143636) | -1.33 |
| | Predicted galacto-N-biose-/lacto-N-biose I ABC transporter, periplasmic substrate-binding protein | 1.31 |
| | Predicted galacto-N-biose-/lacto-N-biose I ABC transporter, permease component 1 | 1.32 |
| Stress response | Gamma-glutamyltranspeptidase PgsD/CapD (EC 2.3.2.2), catalyses PGA anchorage to peptidoglycan | -1.89 |
| | ADA regulatory protein | -1.86 |
| | Cold shock protein CspD | 1.17 |
| | Cold shock protein CspG | 1.21 |
| | Cold shock protein CspB | 1.35 |
| | RNA polymerase sigma factor RpoH | 1.43 |
| | Ribosome-binding factor A | 1.47 |
| | 16 kDa heat shock protein B | 1.49 |
| | 16 kDa heat shock protein A | 1.51 |
| | Alkyl hydroperoxide reductase protein F (EC 1.6.4.-) | 1.59 |
| | Cold-shock DEAD-box protein A | 1.59 |
| | Uptake hydrogenase small subunit precursor (EC 1.12.99.6) | 1.65 |
| | Peptide methionine sulfoxide reductase MsrB (EC 1.8.4.12) | 1.67 |
| | Sigma factor RpoE negative regulatory protein RseB precursor | 1.69 |
| | Non-specific DNA-binding protein Dps | 1.71 |
| | hemimethylated DNA binding protein YccV | 1.74 |
| | Osmotically inducible protein OsmY | 1.80 |
| | RNA polymerase sigma factor RpoS | 1.81 |
| | Cold shock protein CspE | 1.94 |
| | Cold shock protein CspC | 1.96 |
| | Cold shock protein CspA | 2.15 |
| | DedA family inner membrane protein YqjA | 2.18 |
| | SinR, regulator of post-exponential-phase responses genes (competence and sporulation) | -1.93 |
| | Transcriptional regulator of biofilm formation (AraC/XylS family) | -1.93 |

| | | |
|---|---|---|
| Biofilm fromation, adhesion, sensing and competence | Autoinducer 2 (AI-2) ABC transport system, periplasmic AI-2 binding protein LsrB | -1.53 |
| | Type I restriction-modification system, specificity subunit S (EC 3.1.21.3) | -1.52 |
| | Signal peptidase SipW (EC 3.4.21.89), required for TasA secretion | -1.51 |
| | Sporulation kinase B (EC 2.7.13.3) | -1.49 |
| | Internalin-like protein (LPXTG motif) Lmo0409 homolog | -1.48 |
| | Late competence protein ComC, processing protease | -1.46 |
| | Internalin A (LPXTG motif) | -1.43 |
| | Stage V sporulation protein B | -1.41 |
| | RNA polymerase sporulation specific sigma factor SigH | -1.35 |
| | General secretion pathway protein F | -1.35 |
| | MSHA biogenesis protein MshG | -1.32 |
| | Flagellar hook-associated protein FliD | -1.31 |
| | Positive regulator of CheA protein activity (CheW) | -1.30 |
| | Flagellin protein FlaB | -1.26 |
| | Flagellar biosynthesis protein FliC | -1.26 |
| | Flagellin protein FlaA | -1.24 |
| | Internalin D (LPXTG motif) | -1.23 |
| | Flagellar hook subunit protein | -1.22 |
| | Flagellar biosynthesis protein FliQ | -1.22 |
| | Flagellar basal-body rod protein FlgB | -1.20 |
| | Flagellar hook-length control protein FliK | -1.20 |
| | Two-component sensor protein RcsD (EC 2.7.3.-) | 1.32 |
| | FIG002708: Protein SirB1 | 1.35 |
| | Outer membrane protein X precursor | 1.45 |

lues=increased in adult samples
ues= increased in adult samples
